# A membrane-bound nuclease directly cleaves phage DNA during genome injection

**DOI:** 10.1101/2025.11.03.685801

**Authors:** Daniel S. Saxton, Peter C. DeWeirdt, Christopher R. Doering, Ian J. Roney, Michael T. Laub

**Affiliations:** Department of Biology, Massachusetts Institute of Technology, Cambridge, MA 02139, USA; Computational and Systems Biology Program, Massachusetts Institute of Technology, Cambridge, MA 02139, USA; Howard Hughes Medical Institute, Massachusetts Institute of Technology, Cambridge, MA 02139, USA

## Abstract

From mammals to bacteria, the direct recognition and cleavage of viral nucleic acids is a potent defense strategy against viral infection, but it requires mechanisms for distinguishing self from non-self^1,2^. In bacteria, CRISPR-Cas and restriction modification systems achieve this discrimination by recognizing specific DNA sequences or DNA modifications. Alternative mechanisms likely remain to be discovered. Here, we characterize SNIPE, a novel anti-phage defense system that constitutively localizes to the bacterial cell membrane in *E. coli* to block phage λ infection. Using radiolabeled phage DNA and time-lapse microscopy to track phage genomes we demonstrate that SNIPE directly cleaves phage DNA during genome injection.

Based on proximity labeling, we find that SNIPE associates with host proteins essential for λ genome entry and with the λ tape measure protein, which facilitates λ genome injection across the inner membrane. SNIPE also defends against diverse siphoviruses, likely through direct interactions with their tape measure proteins. Our findings establish SNIPE as a widespread bacterial defense system that exploits the spatial organization of phage genome injection to specifically target viral DNA, representing a novel strategy for distinguishing self from non-self in prokaryotic immune systems.

## Introduction

The ability to distinguish self from non-self is a fundamental feature of immune systems across all domains of life. In eukaryotes, pattern recognition receptors (PRRs) enable this distinction by detecting conserved pathogen-associated features, such as lipopolysaccharides (LPS), flagellin, and nucleic acids (e.g., DNA and double-stranded RNA) that are typically absent from the eukaryotic cytoplasm^3–6^. Activated PRRs can trigger various immune responses, including the production of pro-inflammatory cytokines and the initiation of programmed cell death. Similarly, bacteria employ so-called abortive infection systems that often recognize conserved features of invading bacteriophages, such as major capsid proteins, tail proteins, and single-stranded DNA-binding proteins^7–10^. Upon detection, a diverse array of downstream effectors can deplete NAD^+^, degrade nucleic acids, block translation, or permeabilize the cell membrane, among other activities^11–14^. These responses often arrest or kill the host cell, effectively halting phage replication and protecting the surrounding bacterial population.

A second form of non-self recognition involves targeting foreign nucleic acids directly. In eukaryotes, the RNA interference (RNAi) pathway cleaves viral double-stranded RNA into small interfering RNAs (siRNAs)^1,15^ that then guide Argonaute proteins to complementary RNA sequences, leading to the degradation of viral RNA^1,15^. In bacteria, various "direct defense" systems also specifically target foreign DNA. This includes CRISPR-Cas systems and Argonautes that use guide RNAs or DNAs, respectively, to cut foreign DNA in a sequence-specific manner^1,2^. Additionally, restriction modification systems can recognize foreign DNA based on the presence or absence of DNA modifications^16^. Whether additional mechanisms exist for directly identifying and degrading foreign nucleic acids remains an open question.

A recent genetic screen identified a direct defense system, provisionally named PD-λ-1, that is harbored by some strains of *E. coli* and potently blocks phage λ infection^17^. For reasons described below, we re-named this system Surface-associated Nuclease Inhibiting Phage Entry (SNIPE). We demonstrate that SNIPE uses its N-terminal transmembrane domain to localize to the cell membrane, which prevents autoimmune cleavage of host DNA. Using proximity labeling and genetic studies, we show that SNIPE associates with the mannose permease complex in the inner membrane, which is used by λ to promote genome injection. Moreover, proximity labeling during phage genome injection indicates that SNIPE also associates with the phage’s tape measure protein, which is thought to shepherd phage DNA across the inner membrane. Cell biological studies demonstrate that SNIPE cleaves phage DNA during genome injection, thereby preventing phage replication and cell lysis. Thus, our findings suggest that SNIPE provides direct defense by relying on the subcellular localization of phage DNA to distinguish self from non-self.

## SNIPE is a membrane-bound nuclease

To confirm that SNIPE is a direct defense system, we infected cells at a concentration of phage λ such that approximately half of the cells were infected and then monitored cell growth by time-lapse microscopy. In a population of cells lacking SNIPE, infected cells burst and caused a second round of successful phage infection in neighboring cells, as expected (Fig. 1a, Supplemental Movie 1). In sharp contrast, in a population of cells harboring SNIPE, no cell lysis was observed and cells grew to confluence. As a control, we also infected a population of cells containing an abortive infection system, PD-λ-3, and observed that infected cells lysed but prevented a second round of infection^17^. Additionally, growth curve assays showed that SNIPE, but not PD-λ-3, permitted cell survival in the presence of high phage concentrations (Fig. 1b). These results confirm that SNIPE provides direct defense, enabling cells to ward off infection without notably compromising cell growth.

**Figure 1:**
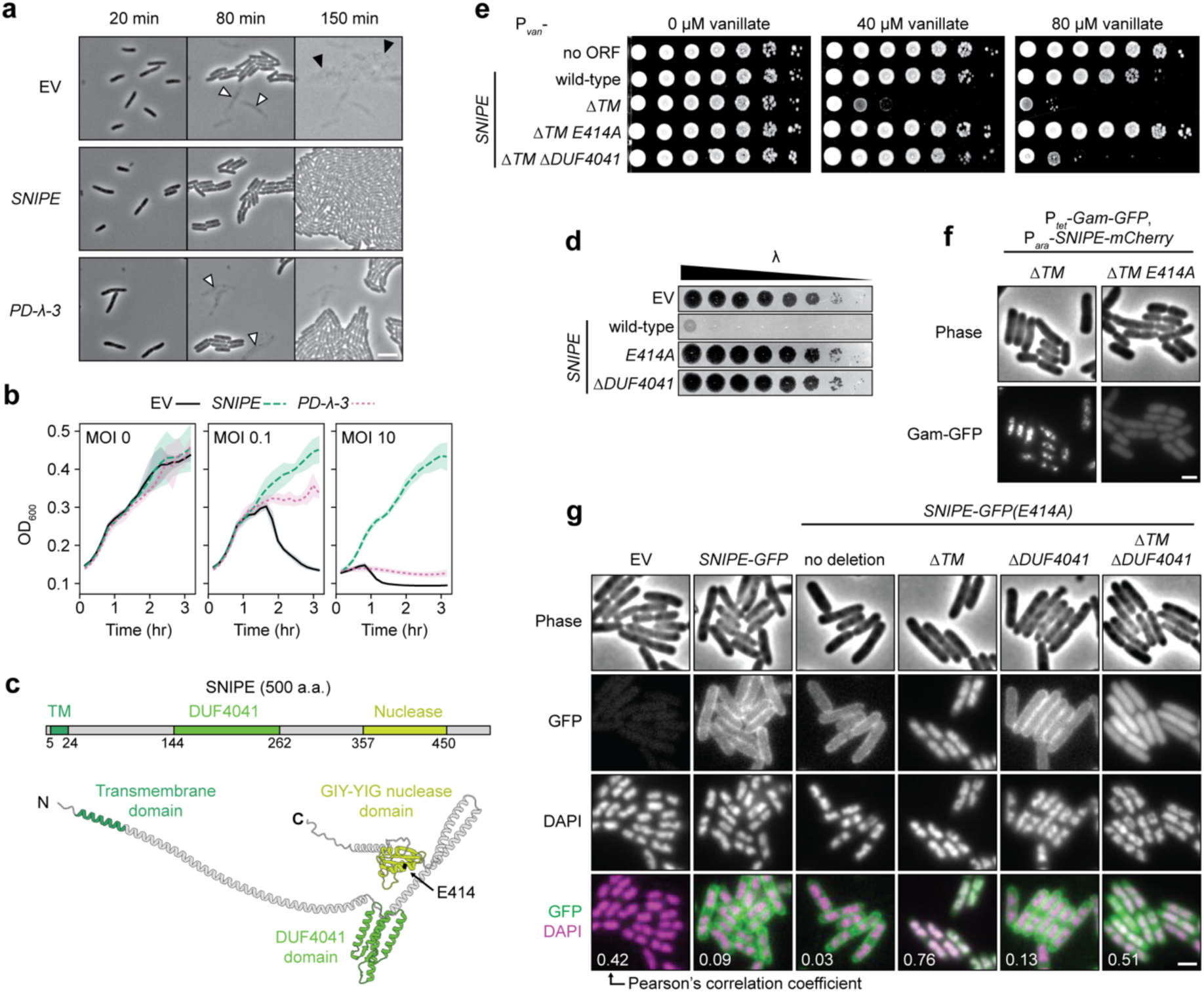
SNIPE is a membrane-bound nuclease that provides direct defense. **(a)** Time course of cells expressing an empty vector (EV), SNIPE, or PD-λ-3. Approximately half of the cells were infected with λ at t = 0 min. White arrows mark cell death from the initial round of infection and black arrows mark cell death from the second round. Scale bar, 3 µm. **(b)** Growth curves for cells expressing an empty vector, SNIPE, or PD-λ-3 and infected at different multiplicities of infection (MOI) of λ at t = 0 min. Line = mean; shaded region = standard deviation. n = 4 independent biological replicates. **(c)** Structure of SNIPE predicted by AlphaFold3, with color-coded domains predicted by HHpred and DeepTMHMM. The N-terminus, C-terminus, and E414A location are indicated. Start and end points of each domain are shown. **(d)** Serial dilutions of λ spotted on lawns of cells expressing an empty vector or different SNIPE constructs. **(e)** Serial dilutions of bacterial strains on plates with 0, 40, or 80 µM vanillate to induce empty vector or P*_van_*-*SNIPE* constructs. **(f)** Single-frame microscopy of cells expressing SNIPE-mCherry constructs and Gam-GFP, a marker of double strand breaks. Scale bar, 1 µm. **(g)** Single-frame microscopy of cells expressing empty vector or SNIPE-GFP constructs. Nucleoids were stained with DAPI. Pearson’s correlation coefficients (R) were calculated for each condition by comparing GFP and DAPI fluorescence signals (n > 100 cells). Scale bar, 1 µm.

To gain insight into the molecular function of SNIPE, we first used a combination of AlphaFold3 and HHpred to predict its domain organization. These analyses suggested that SNIPE is an elongated protein with a single transmembrane domain near the N-terminus, a domain of unknown function (DUF4041) in the middle of the protein, and a GIY-YIG nuclease domain near the C-terminus (Fig. 1c, Extended Data Fig. 1a-b). A deep learning model for transmembrane topology predicted that the N-terminus of SNIPE resides in the periplasm, the single-pass transmembrane domain in the inner membrane, and the rest of the protein in the cytoplasm (Extended Data Fig. 1c)^18^. We tested this model with a membrane topology assay in which a PhoA-LacZα fusion protein was inserted at different locations within SNIPE^19^. PhoA only functions in the periplasm and LacZα only functions in the cytoplasm; thus, the respective activities of these domains indicate the topological orientation of SNIPE. PhoA was only functional when inserted at the N-terminus of SNIPE, and LacZα was only functional when inserted downstream of the transmembrane domain (Extended Data Fig. 1d). These data suggest that SNIPE is anchored in the inner membrane, with the DUF4041 and nuclease domains protruding into the cytoplasm.

To test if the transmembrane, DUF4041, and GIY-YIG nuclease domains are necessary for defense against phage λ, we mutated each of these regions individually and performed plaquing assays. Substituting a predicted catalytic residue in the nuclease domain (E414A) abolished defense, indicating that nuclease activity is essential for SNIPE function (Fig. 1d, Extended Data Fig. 1e). Similarly, a deletion of the DUF4041 ablated defense, suggesting that this globular domain is also critical for SNIPE function. Attempts to clone *SNIPE* lacking the transmembrane domain (*ΔTM*) were unsuccessful, so we put *SNIPE(ΔTM)* under the control of a vanillate inducible (P*_van_*) promoter. Induced expression of this construct was highly toxic, and this toxicity was abolished by the E414A mutation (the toxicity was also partially reduced by ΔDUF4041, which is explored below) (Fig. 1e). These data suggested that SNIPE(ΔTM) may localize to and cleave host DNA. To test this model, we used Gam-GFP, a fluorescently tagged RecBCD inhibitor that localizes to double strand DNA breaks *in vivo*^20^. We first confirmed that this marker was functional by expressing the restriction enzyme EcoRI fused to mCherry, which localized to and cut host DNA, as evidenced by Gam-GFP foci (Extended Data Fig. 1f).

Similarly, we found that SNIPE(ΔTM)-mCherry localized to and cleaved host DNA (Fig. 1f). Consistent with the notion that the E414A mutation disrupts nuclease activity, no double strand breaks were generated by SNIPE(ΔTM E414A)-mCherry.

To assess the subcellular localization of SNIPE and SNIPE(ΔTM) under control of its native promoter, we first fused GFP to the C-terminus of SNIPE and confirmed that this fusion protein retained robust defense against λ infection (Extended Data Fig. 2a). We then found that SNIPE-GFP was expressed and uniformly localized to the cell membrane independent of phage infection (Fig. 1e). To visualize SNIPE(ΔTM)-GFP without the toxic effects of nuclease activity, we used the catalytically inactive variant SNIPE(ΔTM E414A)-GFP. This construct no longer localized to the cell membrane and instead associated with bacterial DNA, as judged by co-localization with the DAPI-stained nucleoid (Fig. 1g, Extended Data Fig. 2b). To confirm these results, we separated cytoplasmic and membrane fractions of cell lysates and found by immunoblotting that SNIPE(E414A)-GFP was strongly enriched in the membrane fraction, whereas SNIPE(ΔTM E414A)-GFP was strongly enriched in the cytoplasm (Extended Data Fig. 2c). These results further suggested that the toxicity of SNIPE(ΔTM) stems from it localizing to and cleaving the host genome. By extension, these data suggest that wild-type SNIPE localizes to the inner membrane to help to sequester its nuclease activity and prevent autoimmunity.

Given that bacterial DNA likely contacts the cell membrane during cellular processes such as chromosome replication and segregation^21^, yet SNIPE is not intrinsically toxic to cells, we hypothesized that membrane-localized SNIPE does not cleave membrane-localized host DNA. To test this, we ectopically localized host DNA to the cell membrane by fusing two transmembrane domains from MalF to GFP-Fis, a fluorescently tagged DNA-binding protein. Strong expression of MalF(TM1-2)-GFP-Fis localized DAPI-stained DNA to the cell membrane and was highly toxic even in the absence of SNIPE (Extended Data Fig. 2d). Across a range of MalF(TM1-2)-GFP-Fis expression levels with varying degrees of toxicity, the presence of SNIPE generated no additional toxicity for cells. This result indicates that localization of host DNA to the cell membrane does not make it susceptible to SNIPE-mediated cleavage, suggesting that SNIPE exists in an auto-inhibited state when localized to the membrane.

To understand why removal of the DUF4041 reduced the toxicity of SNIPE(ΔTM) (Fig. 1e), we examined the electrostatic surfaces of the AlphaFold-predicted SNIPE structure and found that the DUF4041 domain contains a positively charged surface that may facilitate DNA binding (Extended Data Fig. 2e). To test this hypothesis, we compared the localization of SNIPE(ΔTM E414A)-GFP with and without the DUF4041 domain. Indeed, whereas SNIPE(ΔTM E414A)-GFP localized to the nucleoid, SNIPE(ΔTM E414A ΔDUF4041)-GFP exhibited diffuse cytoplasmic localization (Fig. 1e). Additionally, we found that a version of SNIPE containing only the DUF4041 and the downstream alpha helix fused to GFP localized to the bacterial nucleoid (Extended Data Fig. 2f). These observations suggest that the DUF4041 promotes DNA binding by SNIPE(ΔTM). Collectively, our results support a model in which SNIPE contains a functional nuclease domain, a DUF4041 domain that facilitates DNA binding, and a transmembrane domain that anchors SNIPE to the inner membrane, preventing autoimmune cleavage of host DNA.

## SNIPE cuts phage DNA during genome injection

Given that SNIPE localizes to the membrane and contains a functional nuclease, we reasoned that SNIPE may directly cleave phage DNA during genome injection. We first confirmed that SNIPE does not affect adsorption of phage λ (Fig. 2a), consistent with a previous study^17^. Next, we sought to directly visualize genome injection using a fluorescence assay in which λ carries a *parS* site and *E. coli* produces CFP-ParB^22^. After phage genome injection, CFP-ParB oligomerizes on *parS* sites and λ*^parS^* genomes appear as fluorescent puncta. Consistent with previous work^22^, CFP-ParB puncta appeared within ten minutes of λ*^parS^* infection and expanded during the course of phage genome replication, which was followed by cell lysis (Fig. 2b-c, Supplemental Movie 2). In contrast, the number of CFP-ParB puncta that appeared in SNIPE-expressing cells was reduced by approximately 30-fold, and there was a concomitant reduction in cell lysis. In rare cases where a λ*^parS^* genome appeared in a SNIPE-containing cell, it went on to replicate and lyse the cell, suggesting that if phages stochastically evade SNIPE at the cell membrane their development proceeds unimpeded (Extended Data Fig. 3a-b, Supplemental Movie 2). The SNIPE-dependent reduction in CFP-ParB puncta was not observed with the catalytically inactive E414A mutation, indicating that this reduction was dependent on nuclease activity (Fig. 2b-c).

**Figure 2:**
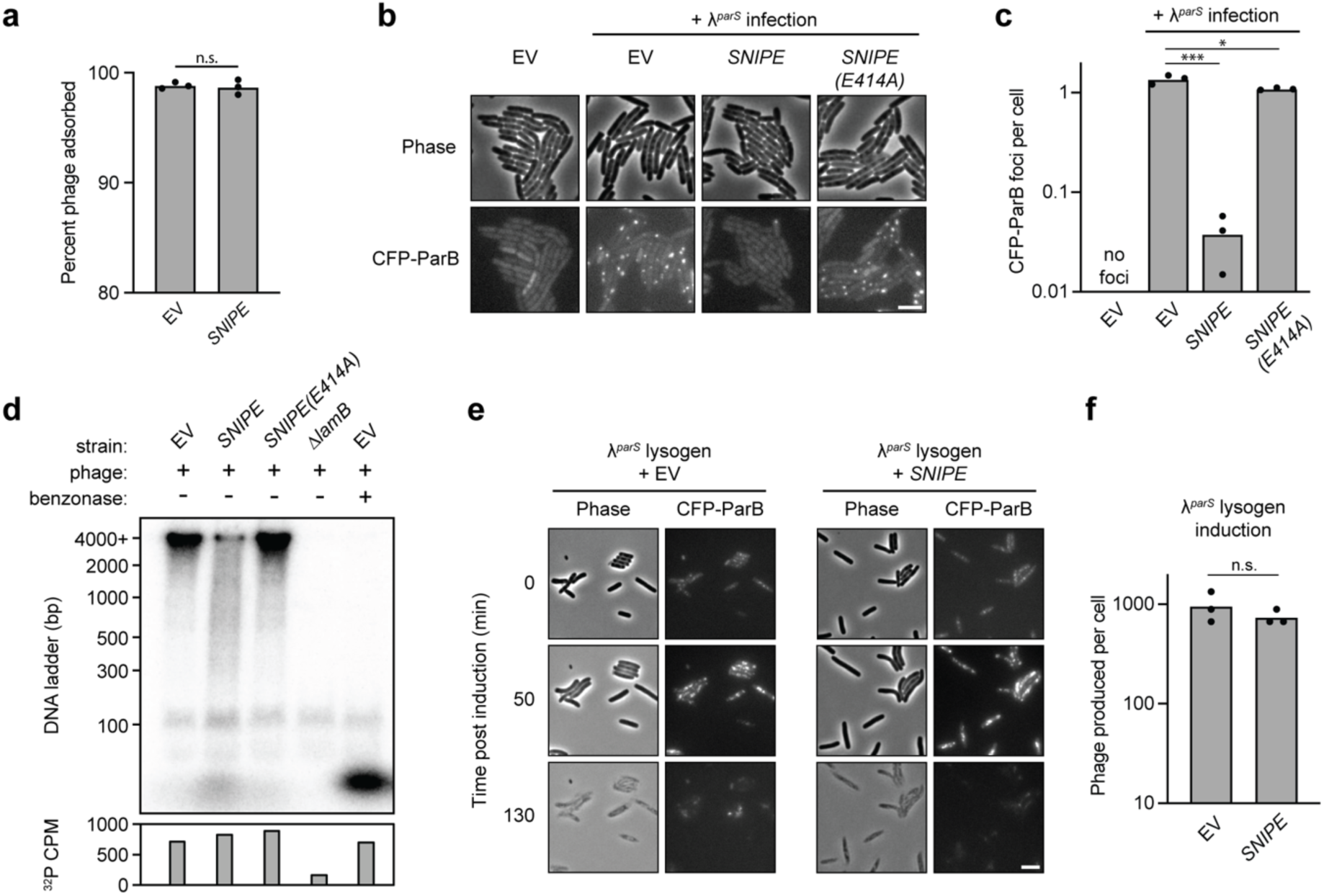
SNIPE cleaves phage DNA during genome injection. **(a)** Adsorption assay with λ and cells containing SNIPE or an empty vector. Summary of 3 independent replicates. n.s. indicates p = 0.76 (unpaired two-tailed t-test). **(b)** Microscopy of cells expressing CFP-ParB and an empty vector or different SNIPE constructs. Cells were infected with λ^parS^ and imaged 5 min after genome injection. Scale bar, 2 µm. **(c)** Quantification of CFP-ParB foci per cell from (**b)** and two additional independent replicates. * indicates p = 0.03 and *** indicates p < 0.0001 (unpaired two-tailed t-tests). **(d)** Different bacterial strains were infected with ^32^P-labeled λ and lysed 15 minutes after genome injection. Benzonase was added to one lysate sample. Lysates were subjected to electrophoresis through a polyacrylamide gel and imaged with a phosphorscreen. Total ^32^P per sample was measured with a scintillation counter. **(e)** λ*^parS^* lysogens expressing the heat-labile cI857 repressor, CFP-ParB, and SNIPE or an empty vector were induced via heat shock at 42 °C. Time-lapse microscopy was performed to monitor CFP-ParB foci dynamics and cell lysis. Scale bar, 2 µm. **(f)** Quantification of plaque forming units (PFU) per induced cell was performed for strains in (**e)** and two additional independent replicates. n.s. indicates p = 0.35 (unpaired two-tailed t-test).

Although these experiments were consistent with SNIPE cleaving incoming phage DNA, they did not formally rule out the possibility that SNIPE permits phage adsorption but somehow blocks genome injection. To directly test if phage DNA is cleaved by SNIPE, we adapted the classic Hershey-Chase experiment^23^. In this adaptation, we infected cells with λ containing ^32^P-labeled DNA, harvested cells shortly after genome injection, and then measured the size of radiolabeled DNA fragments. Infection of empty vector cells yielded a clear ^32^P-labeled band at the upper limit of size detection (4000 bp), as would be expected for the injected, ∼42,000 bp λ genome (Fig. 2d). This band was cleaved into a <100 bp band upon benzonase treatment, and this smaller band likely represents mononucleotides as [γ-^32^P]-ATP formed a band at the same size (Extended Data Fig. 3c). Consistent with the ^32^P-labeled band originating from injected phage DNA, it was not observed upon infection of cells lacking the λ receptor (Δ*lamB*). In contrast to empty vector cells, infection of SNIPE-containing cells yielded a smear of DNA fragment sizes ranging from >4000 bp to <100 bp, as well as the band that likely corresponded to mononucleotides. This profile of ^32^P-labeled DNA cleavage was reversed in cells expressing SNIPE(E414A). Importantly, the amount of injected ^32^P was similar across empty vector, SNIPE, and SNIPE(E414A)-expressing cells, as measured by scintillation counting (Fig. 2d, Extended Data Fig. 3c). Taken together, our results strongly suggest that SNIPE does not block phage genome injection, but rather cleaves phage DNA during the injection process.

To provide an orthogonal test of our model, we asked if SNIPE could target a phage genome that was already in the cell. To this end, we introduced a plasmid expressing SNIPE or carrying an empty vector into a λ*^parS^* lysogen that produces CFP-ParB and encodes a temperature-sensitive cI repressor, which causes the λ*^parS^*prophage to enter the lytic cycle at high temperatures. Upon heat shock, λ*^parS^*prophages were induced and then replicated, as manifest by CFP-ParB foci, followed by synchronous cell lysis (Fig. 2e, Supplemental Movie 3). As predicted, SNIPE did not qualitatively affect the dynamics of CFP-ParB foci or cell lysis, and it did not significantly affect the number of phage particles produced by prophage induction (Fig. 2f). These findings support the model that SNIPE cleaves phage DNA during, but not after, genome injection.

## SNIPE interacts with the mannose permease complex

Collectively, our results indicated that SNIPE localizes to phage genome injection sites where it can cleave incoming DNA. But how does SNIPE find such sites? After adsorbing to the outer membrane receptor LamB, phage λ requires the inner membrane components of the mannose permease complex (consisting of ManY and ManZ) for genome injection, though molecular details of this process remain poorly understood^24,25^. In principle, SNIPE could target phage DNA if it associates with LamB or ManYZ. To test these possibilities, we first turned to a "generalist" mutant of λ that evolved to infect *ΔlamB* and *ΔmanYZ* strains^26^. This phage has a mutation in the tail tip protein that allows it to bind either LamB or an alternate outer membrane protein, OmpF (Fig. 3a). This phage also has a mutation in the tape measure protein that allows it to circumvent a requirement for ManYZ, though the nature of this alternate genome injection pathway remains unclear.

**Figure 3:**
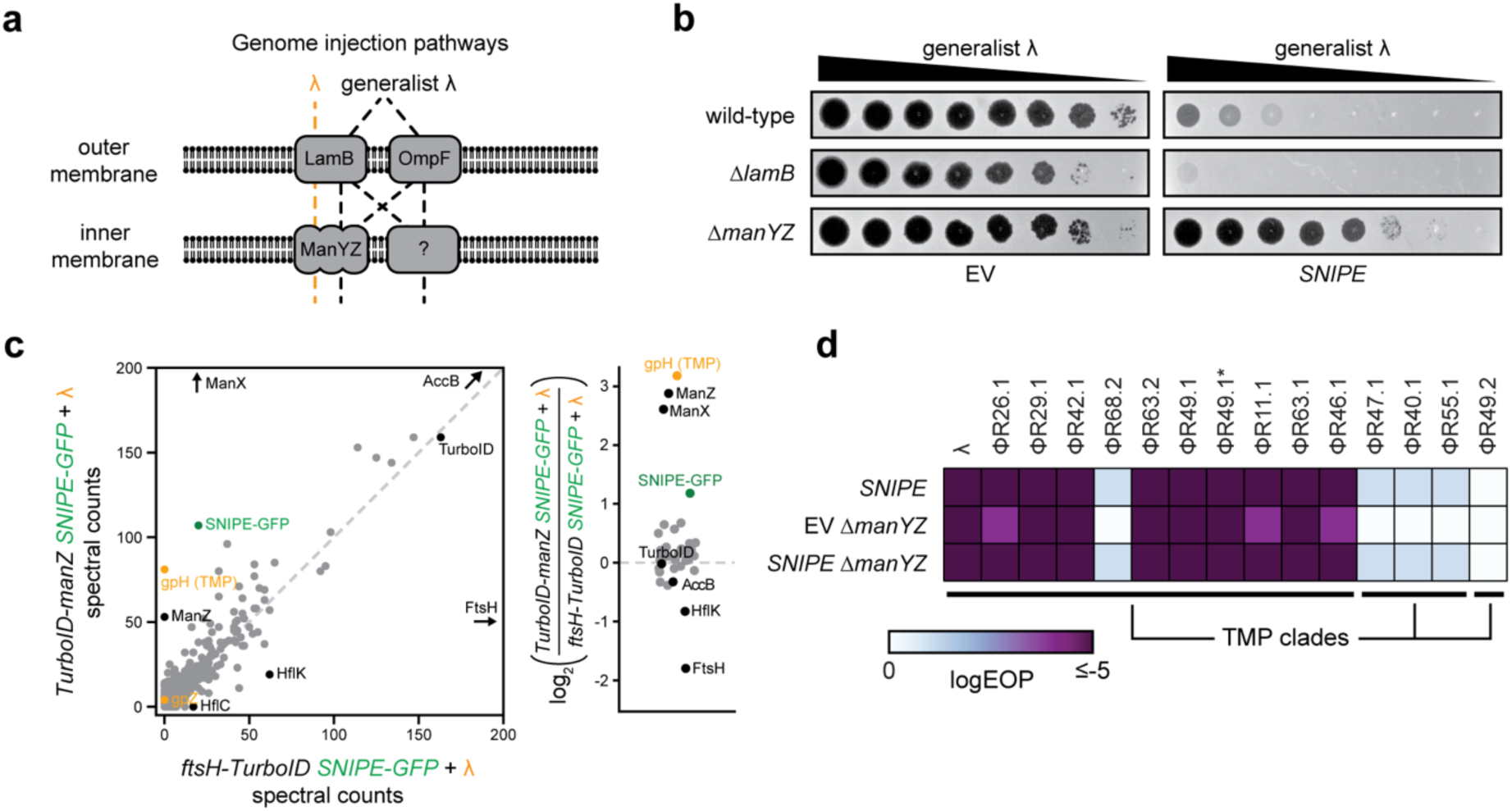
SNIPE interacts with ManYZ. **(a)** Schematic of genome injection routes used by λ and the generalist λ mutant^26^. **(b)** Serial dilutions of the generalist λ mutant were spotted onto bacterial lawns with different genomic deletions and an empty vector or SNIPE-expressing plasmid. **(c)** Proximity labeling was performed with infected cells expressing SNIPE-GFP and TurboID-ManZ or FtsH-TurboID. Biotinylated proteins were enriched with streptavidin beads and quantified by mass spectrometry. Log_2_ ratios of spectral counts for proteins with ≥ 50 spectral counts in at least one sample are shown on the right. A pseudocount was added to each spectral count value to facilitate ratio calculations. Phage proteins are labeled in orange, SNIPE-GFP is labeled in green, and other proteins of interest are labeled in black. **(d)** Efficiency of plaquing (EOP) data for λ and other temperate phages on wild type or *ΔmanYZ* cells expressing SNIPE or harboring an empty vector. Phages in a given TMP clade share > 90% amino acid identity between their TMPs, and < 20% identity with TMPs outside of the clade.

We first confirmed that the generalist λ mutant was able to infect *ΔlamB* and *ΔmanYZ* strains, albeit with minor plaquing defects (Fig. 3b). Next, we reasoned that if SNIPE-mediated defense requires LamB or ManYZ, then shunting genome injection through one of the alternate pathways would circumvent defense. SNIPE provided robust defense against the generalist λ mutant in both wild-type and *ΔlamB* cells, demonstrating that LamB is not required for defense. In contrast, SNIPE-mediated defense was strongly reduced in *ΔmanYZ*. These data suggest that although the generalist λ mutant can use various genome injection pathways, ManYZ is the preferred pathway through the inner membrane if provided. As such, the generalist λ mutant likely uses ManYZ in wild-type and *ΔlamB* cells, and is susceptible to SNIPE, while forcing this phage to use an alternate pathway in *ΔmanYZ* cells largely circumvents SNIPE defense.

Our genetic analysis suggested that SNIPE requires the mannose permease complex to provide robust defense against λ. To more directly assess interactions between SNIPE and ManYZ, we turned to biotin ligase proximity labeling. This method involves fusing TurboID, which generates diffusible 5’-biotinyl-AMP that conjugates with nearby lysines, to a protein of interest and identifying biotinylated proteins via streptavidin pulldown and mass spectrometry^27^. We first generated SNIPE-TurboID but found it did not provide strong defense in the presence of biotin (Extended Data Fig. 4a). We therefore generated a plasmid expressing the *manXYZ* operon containing *TurboID-manZ*. This construct complemented λ plaquing defects on a *ΔmanXYZ* strain and did not affect SNIPE-mediated defense, indicating that the TurboID-ManZ fusion was functional (Extended Data Fig. 4b).

To perform proximity labeling during phage genome injection, we added wild-type λ and exogenous biotin to cells producing ManXY, TurboID-ManZ, and SNIPE-GFP. At 15 minutes post-infection, we lysed cells and used streptavidin pulldowns to isolate biotinylated proteins, which were identified by mass spectrometry. As expected, one of the most enriched proteins was AccB, the only protein that is naturally biotinylated in *E. coli* (Fig. 3c, Extended Data Fig. 4d). We also observed strong enrichment of ManX, a known interaction partner of ManYZ, providing a proof of concept for this assay; ManY was not enriched, but only harbors one cytoplasmic lysine that could be biotinylated^28^. Notably, SNIPE-GFP was also enriched, and the only phage protein that was robustly labeled by TurboID-ManZ during genome injection was the λ tape measure protein (TMP), also termed gpH (additional tail components gpJ and gpZ were weakly labeled in some TurboID-ManZ samples). To assess the specificity of these interactions, we fused TurboID to one of two other inner membrane proteins, ProW or FtsH, and again performed proximity labeling during λ infection. We found that ManX, SNIPE-GFP, and the λ tape measure protein were enriched only in the TurboID-ManZ samples (Fig. 3c, Extended Data Fig. 4c-e).

These results suggest that ManYZ specifically associates with SNIPE and the λ tape measure protein during genome injection.

To test whether SNIPE interacts with ManYZ independent of the tape measure protein, we performed proximity labeling with TurboID-ManZ in the presence and absence of phage infection. Indeed, we observed strong enrichment of SNIPE-GFP both in the presence and absence of phage infection, and similar results were obtained in a similar experiment involving untagged SNIPE (Extended Data Fig. 4f-g). These observations support the notion that SNIPE and ManYZ interact *in vivo* prior to and during λ infection. Notably, this interaction did not impair ManYZ-mediated transport of mannose across the inner membrane, as SNIPE-containing cells were able to metabolize mannose on MacConkey agar (Extended Data Fig. 4h). We also asked whether the tape measure protein interacts with ManYZ independently of SNIPE. During genome injection, we observed that TurboID-ManZ labeled the tape measure protein at similar levels in the presence and absence of SNIPE (Extended Data Fig. 4i). Together, these data suggest that ManYZ interacts with both SNIPE and the tape measure protein, independent of each other. By extension, our results indicate that SNIPE interacts with ManYZ prior to infection, positioning it to also associate with the tape measure protein during infection and thereby target the incoming phage DNA for degradation.

If SNIPE interacts with ManYZ, then it should defend against other phages that also use the ManYZ complex for genome injection. To test this prediction, we screened a panel of temperate phages that are relatives of λ^29^. Most of these phages were not able to infect a Δ*manYZ* strain, suggesting that they use ManYZ for genome injection (Fig. 3d). These same phages were highly susceptible to SNIPE-mediated defense. In contrast, phages that did not require ManYZ for infection were either unaffected or only modestly inhibited by SNIPE. This modest inhibition still occurred in *SNIPE ΔmanYZ* cells, indicating that SNIPE offers weak defense independent of ManYZ, an idea explored further below. Overall, the correlation between SNIPE susceptibility and use of ManYZ supports our model that SNIPE interacts with ManYZ to provide robust defense against phages that use this protein complex for genome injection.

Given our TurboID analyses, we also generated an alignment of the tape measure proteins encoded by each of the phages in this panel. These tape measure proteins formed three distinct clades that strongly correlated with susceptibility to SNIPE and dependence on ManYZ (Fig. 3d, Extended Data Fig. 5a). Interestingly, in the clade that was highly susceptible to SNIPE and that required ManYZ for infection there was one exception, the phage ΦR68.2. When comparing the ΦR68.2 tape measure protein to other tape measure proteins in this clade, it harbored a region from amino acids 300-400 that differed substantially from the others (Extended Data Fig. 5b).

Notably, this region overlaps with the A304V mutation in the tape measure protein of the generalist λ mutant that allows it to infect *ΔmanYZ* cells^26^. These results support the notion that phage genome injection via ManYZ substantially increases its susceptibility to SNIPE. However, SNIPE can provide some defense independent of ManYZ, as evidenced by the modest protection against phages ΦR40.1, ΦR47.1, ΦR55.1,and ΦR68.2 and by the reduced, but not fully abolished, defense that SNIPE provides against the "generalist" λ mutant in the *ΔmanYZ* background.

## SNIPE can target siphoviruses independent of ManYZ

To further test the molecular determinants of SNIPE-mediated defense and its phage specificity, we screened the BASEL collection of virulent phages for their susceptibility to SNIPE and their dependence on ManYZ^30^. None of the myoviruses or podoviruses in the BASEL collection were susceptible to SNIPE defense (Fig. 4a). However, the large majority of siphoviruses were weakly targeted by SNIPE, with a subset targeted more robustly. Surprisingly, none of these phages showed plaquing defects on *ΔmanYZ*, and SNIPE-mediated defense was unaffected in *ΔmanYZ* cells. These data indicate that SNIPE offers broad defense against most siphoviruses independent of ManYZ.

**Figure 4:**
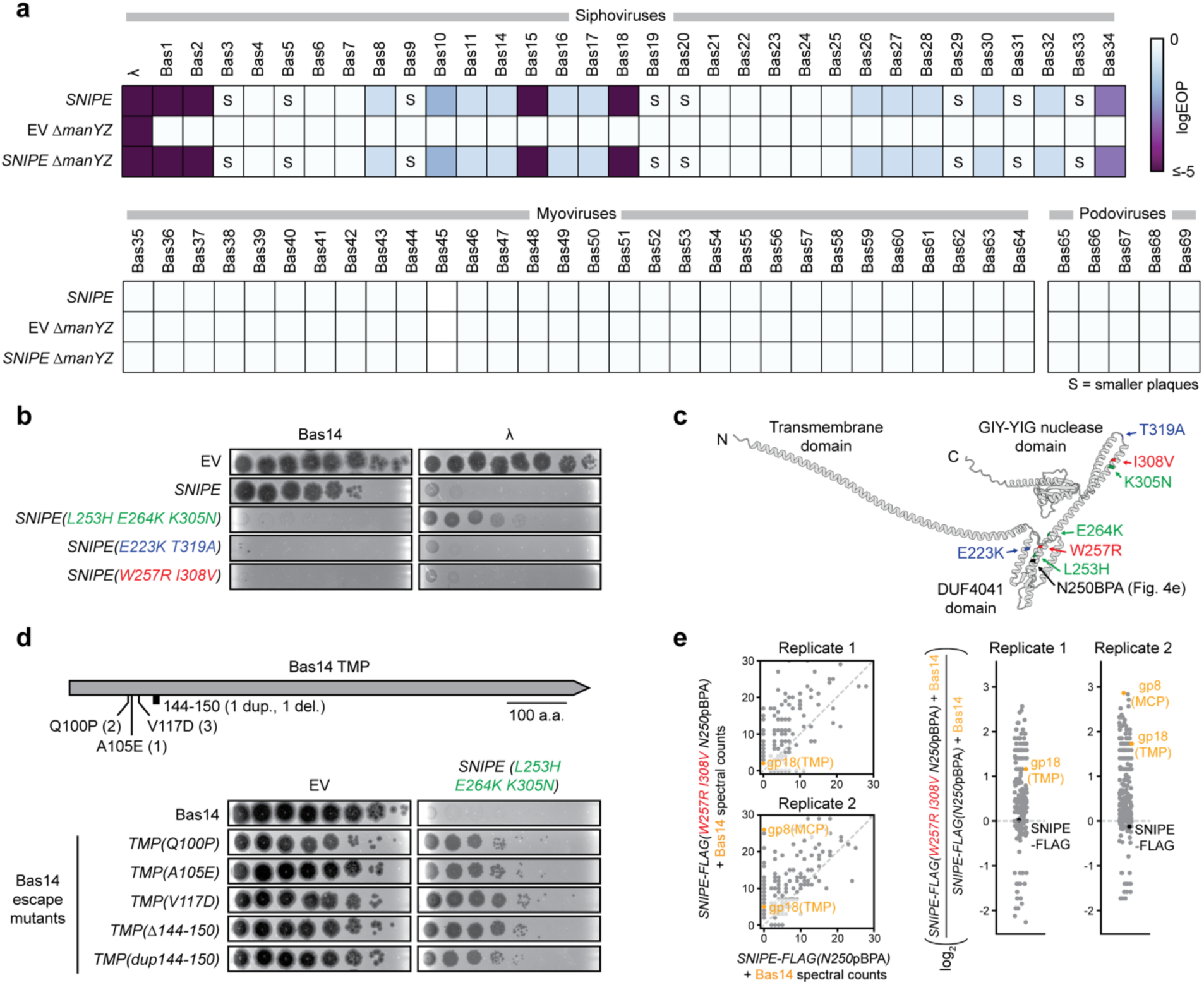
SNIPE can target siphoviruses independent of ManYZ. **(a)** EOP data for λ and BASEL phages on wild type or *ΔmanYZ* cells expressing SNIPE or harboring an empty vector. Smaller plaque sizes are indicated with an ’S’. **(b)** Bas14 and λ were spotted onto lawns of *E. coli* expressing an empty vector, SNIPE, or the indicated SNIPE mutants. **(c)** Mutations that enhance SNIPE-mediated defense against Bas14 are highlighted on the structure of SNIPE predicted by AlphaFold3. The N250 residue replaced with pBPA in Fig. 4e is also indicated. **(d)** Summary of identified escape mutations, all of which map to the Bas14 TMP. The number of independently isolated escaper plaques with a given genotype is shown in parentheses. Bas14 and Bas14 escape mutants were spotted onto lawns of empty vector and SNIPE(L253H E264K K305N)-expressing cells. **(e)** UV-induced crosslinking of Bas14-infected cells expressing SNIPE-FLAG(W257R I308V N250pBPA) or SNIPE-FLAG(N250pBPA). These proteins and crosslinked products were pulled down with anti-FLAG beads and quantified by mass spectrometry. Scatterplot data are zoomed in to visualize proteins with low spectral counts; unzoomed data are shown in Extended Data Fig. 8c. Log_2_ ratios of spectral counts are shown on the right. A pseudocount of 0.5 was added to each spectral count value to facilitate ratio calculations. Phage proteins are labeled in orange, and SNIPE-FLAG is labeled in black.

As noted above, SNIPE-TurboID was non-functional and thus could not be used to examine its interactions with proteins from these other phages. Thus, to probe the basis of SNIPE’s broad defense against siphoviruses, we used error-prone PCR to create a pool of cells harboring mutagenized SNIPE and challenged it with phage Bas14. While wild-type SNIPE provided approximately 1 log-fold protection against Bas14, three isolated SNIPE mutants strongly enhanced this defense, with each offering 6- to 7-log fold protection (Fig. 4b). Each of these SNIPE mutants had at least two point mutations that mapped to the DUF4041 and the downstream alpha helix (Fig. 4c). Interestingly, cloning single mutations *de novo* revealed that each mutation identified in the DUF4041 domain was sufficient to strongly enhance defense against Bas14, whereas mutations in the downstream alpha helix only partially enhanced this defense (Extended Data Fig. 6a-b). We suspect that these mutations were recovered together due to the extreme selective pressure imposed by Bas14 infection in the original screening step.

Given that Bas14 does not require ManYZ for entry, we inferred that these mutations in the DUF4041 domain enhanced the ManYZ-independent mechanism employed by SNIPE.

To test if the SNIPE mutants we isolated had improved defense generally or only against specific phages, we screened these mutants against λ and the siphoviruses in the BASEL collection. All three SNIPE mutants enhanced defense only against Bas14-18, which constitutes a clade of highly related phages (Extended Data Fig. 6c). This finding suggested that these SNIPE mutants enhance binding to part of the genome injection apparatus used by Bas14-18. To identify the basis of this enhanced defense, we selected Bas14 escapers on the SNIPE mutant harboring the substitutions L253H, E264K, and K305N. Strikingly, the only mutations found in escape phages mapped to the Bas14 gene that encodes the tape measure protein (Fig. 4d). Three of these mutations were substitutions (Q100P, A105E, and V117D), one was a deletion of amino acids 144-150, and one was a duplication of amino acids 144-150. All of the escape mutants also provided some degree of escape against the other SNIPE mutants (E223K T319A and W257R I308V), though they were slightly more susceptible to wild-type SNIPE for reasons that remain unclear (Extended Data Fig. 7a-b). Notably, Bas14-Bas18 have tape measure proteins that are >90% identical to each other and <20% identical to any other tape measure protein in the collection (Extended Data Fig. 6c), and the region of the tape measure protein that was mutated in Bas14 escapers (amino acids 100-150) had >85% sequence identity only within the Bas14-18 clade (Extended Data Fig. 7c). Thus, Bas14 can escape the robust defense of SNIPE mutants by mutating a conserved region of its tape measure protein.

How do these SNIPE mutants provide enhanced defense in a manner that is dependent on the tape measure protein? One possibility is that SNIPE binds weakly to diverse siphovirus tape measure proteins, and the SNIPE mutants strengthen this interaction for phages in the Bas14-18 clade. Alternatively, the SNIPE mutants may enhance binding to an inner membrane protein used by the Bas14-18 clade, and Bas14 escaper mutations may switch to a using a different inner membrane protein. To test the latter possibility, we performed transposon-insertion sequencing (Tn-Seq)^31^. Specifically, we used λ, Bas14, or the Bas14 *TMP(A105E)* escape isolate to infect pools of cells in which barcoded transposons are inserted throughout the genome, such that most cells will die from infection, but cells harboring a transposon in a gene necessary for phage infection will survive. As expected, this screen identified ManY and ManZ as required for infection of λ (Extended Data Fig. 8a). In contrast, no integral inner membrane proteins were required for infection of either Bas14 or Bas14 *TMP(A105E)*. Notably, one or both of these phages may require essential inner membrane proteins, which cannot be identified with Tn-Seq. Nevertheless, we find it unlikely that relatively minor mutations in Bas14 escapers conferred a complete switch from one essential inner membrane protein to another, and instead favor a model in which these phages do not use a specific host inner membrane protein for genome injection. By extension, our results indicate that the SNIPE mutants may enhance binding to the Bas14 tape measure protein, which is disrupted in Bas14 escapers.

To test if the SNIPE DUF4041 physically interacts with the Bas14 tape measure protein, we turned to crosslinking with the unnatural amino acid *p-*benzoylphenylalanine (pBPA). In this approach, we replaced the SNIPE N250 codon with an amber codon and expressed a specialized tRNA synthetase that can incorporate pBPA at this site^32^. Given that N250 is adjacent to the W257R mutation that enhanced defense against Bas14, we reasoned pBPA incorporated at this site may crosslink to the Bas14 tape measure protein (Fig. 4c). First, we confirmed that SNIPE-FLAG(W257R I308V N250pBPA) was still able to defend against Bas14 (Extended Data Fig. 8b). Next, we infected this strain with Bas14, exposed cells to UV to crosslink pBPA to nearby proteins, and used anti-FLAG pulldowns to identify crosslinked proteins with mass spectrometry. The only phage protein recovered across both replicates was the tape measure protein (Fig. 4e, Extended Data Fig. 8c-d). In contrast, no phage proteins were detected under similar conditions with SNIPE-FLAG(N250pBPA), suggesting that recovery of phage proteins was dependent on the W257R I308V mutations. Together, these results indicate that the W257R I308V mutations enhance binding to the Bas14 tape measure protein. Therefore, our results support a model in which wild-type SNIPE provides broad defense against siphoviruses by weakly interacting with siphovirus tape measure proteins. This defense could then be augmented by SNIPE binding to an inner membrane protein used by the phage, like ManYZ as with λ, or by a strengthened interaction between the DUF4041 and a specific type of tape measure protein, as with Bas14.

## SNIPE homologs have multiple diversified domains

To elucidate how evolutionary pressures shape the functional domains of SNIPE, we analyzed a set of previously identified SNIPE homologs^17^. Consistent with this prior study, 33% of well-sequenced bacterial clades harbor at least one SNIPE homolog and several clades harbor SNIPE homologs in up to 10% of their genomes, indicating that SNIPE provides anti-phage defense in diverse species (Extended Data Fig. 9). We curated this set of SNIPE homologs with the clustering algorithm MMseqs2 at 95% identity to remove highly similar sequences and better understand the sequence diversity these homologs^33^; this clustering yielded a set of ∼500 homologs distributed across many bacterial phyla (Fig. 5a). The most highly conserved region of SNIPE was the GIY-YIG nuclease domain, supporting the notion that nuclease activity is a critical feature of SNIPE (Fig. 5b). In contrast, the DUF4041 domain was less well conserved, and the N-terminal region had very low sequence conservation. When quantifying the length of each domain in each homolog, we additionally found that the N-terminal region had highly variable lengths compared to the other regions (Fig. 5c). The diversity of the N-terminal region may suggest that it functions as an "adapter" that plays a role in determining the phage specificity of SNIPE homologs.

**Figure 5:**
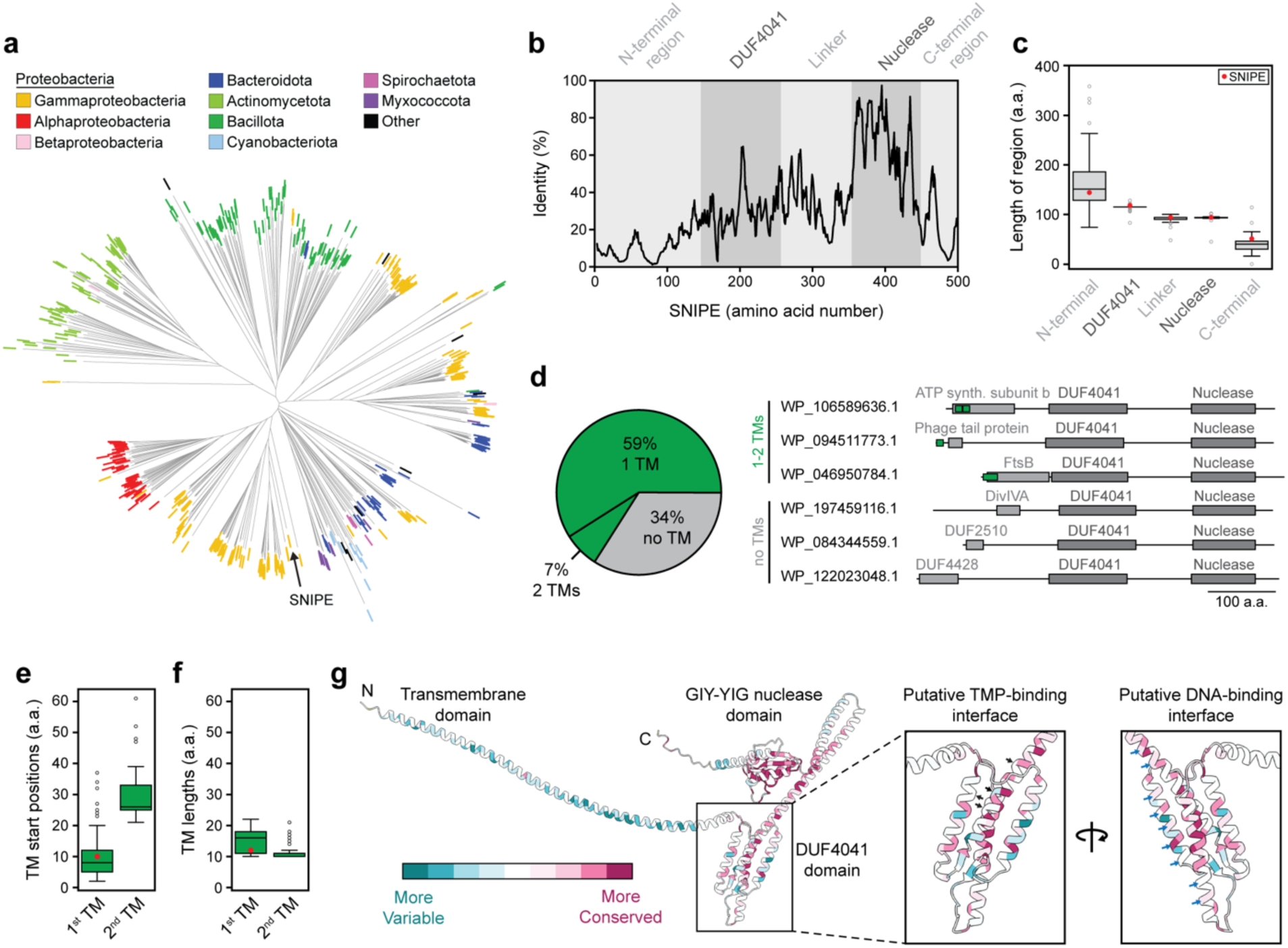
SNIPE homologs harbor diverse N-terminal regions. **(a)** Unrooted phylogenetic tree of 474 SNIPE homologs with tip colors representing bacterial classes (alpha-, beta-, and gamma-proteobacteria) or phyla (all other categories) that harbor these homologs. **(b)** Sequence identity plot of SNIPE homologs, aligned and numbered relative to the amino acid positions of SNIPE. A 5-residue rolling mean was applied. **(c)** Length distributions of the extracted homologous sequences for each region of interest. The length of each region of SNIPE is shown as a red dot for reference. Center line, median; box limits, upper and lower quartiles; whiskers, 1.5x interquartile range; points, outliers. n = 474 SNIPE homologs. **(d)** Percentages of SNIPE homologs that contain 0, 1, or 2 transmembrane domains, as predicted by DeepTMHMM. Representative examples of SNIPE homologs with distinct N-terminal regions, predicted by HMMER, are shown as schematics. **(e)** Distributions of start positions for predicted transmembrane domains of SNIPE homologs. The length of the SNIPE transmembrane domain is labeled as a red dot for reference. Center line, median; box limits, upper and lower quartiles; whiskers, 1.5x interquartile range; points, outliers. n = 313 1^st^ TM domains, n = 33 2^nd^ TM domains. **(f)** Same as in (**e)**, but for transmembrane domain lengths. **(g)** Evolutionary conservation of amino acids as estimated by ConSurf, overlayed onto the predicted structure of SNIPE. Insets indicate different interfaces of the DUF4041. Black arrows indicate locations of SNIPE mutations that enhanced defense against Bas14. Blue arrows indicate positively charged amino acids that may faciliate DNA binding.

Given that the N-terminal transmembrane domain of *E. coli* SNIPE is critical for its function (Fig. 1e), we next asked whether membrane localization is common among the N-terminal regions of SNIPE despite their lack of sequence similarity. Indeed, a deep learning model for transmembrane domains predicted that 59% of SNIPE homologs harbored one transmembrane domain and 7% harbored two transmembrane domains (Fig. 5d)^34^. These transmembrane domains were predicted to reside within the first 70 amino acids of SNIPE homologs and ranged from 10-20 amino acids in length (Fig. 5e-f). HMMER analysis of these SNIPE homologs containing predicted transmembrane domains indicated that their N-terminal regions are often similar to known bacterial inner membrane proteins, such as FtsB, PilO, and PspB (Fig. 5d, Extended Data Fig. 10a). Additionally, some N-terminal regions were homologs of the phage P22 tail needle protein, which is implicated in cell envelope penetration^35^. Thus, despite the low sequence conservation of N-terminal regions, most of these regions likely target SNIPE homologs to the cell membrane and may interact with host and/or phage proteins to enhance defense.

For the remaining 34% of SNIPE homologs that did not contain a predicted TM, we asked whether the N-terminal regions of these homologs contain other features that may target them to the membrane. A combination of AlphaFold2, structural homology searches, and sequence homology searches with HMMER indicated that SNIPE homologs lacking predicted transmembrane domains frequently harbor domains from proteins such as DivIVA and type III secretion system ATPases, which localize to bacterial membranes by recognizing high negative membrane curvature and binding to inner membrane proteins, respectively (Fig. 5d, Extended Data Fig. 10a-b). Strikingly, some of these SNIPE homologs are predicted to harbor N-terminal globular domains with structural and sequence homology to Pleckstrin Homology (PH) domains, which can localize to membranes by binding to phospholipids (Extended Data Fig. 10b-f).

Together, our analyses indicate that SNIPE homologs can localize to membranes by mimicking phage or bacterial proteins that harbor transmembrane domains or associate with membranes by other means.

Another critical feature of SNIPE is the DUF4041 domain, which is required for defense, promotes DNA binding, and likely interacts with a conserved feature of siphovirus tape measure proteins. To investigate DUF4041 evolution, we used ConSurf, an algorithm that measures the diversity of amino acid sequences across homologs of a protein of interest and maps those diversity scores onto the protein structure^36^. These analyses showed that the positively charged region of the DUF4041 domain is conserved relative to the rest of the domain, and that SNIPE homologs usually maintained positively charged residues in this region (Fig. 5g, Extended Data Fig. 10g). These data further support the notion that the positively charged interface binds DNA and that this binding activity is broadly conserved. In contrast, the interface of the DUF4041 that likely binds to tape measure proteins exhibited more variability (Fig. 5g). In particular, the E223K and W257R mutations that were sufficient to enhance defense against Bas14-18 are frequently found in SNIPE homologs (Extended Data Fig. 10g). These results are consistent with a model in which the DUF4041 has been moderately diversified to strengthen binding to specific types of tape measure proteins. Taken together, our results indicate that the N-terminal region and the DUF4041 of SNIPE are under selective pressure to bind to distinct host and/or phage proteins to target different phages during genome injection.

## Discussion

Computational and genetic screens continue to reveal new anti-phage defense systems at a rapid pace^17,37–39^. Many of these newly discovered systems function as abortive infection systems in which the host cell dies or arrests growth to prevent spread of a phage through the population. In contrast, direct defense systems enable an infected cell to ward off the infection and continue proliferating. Such systems require the ability to discriminate between self and non-self so that the host is spared while the virus is targeted. Most famously, CRISPR-Cas and restriction modification systems identify non-self DNA based on sequence specificity and DNA modifications, respectively^40,41^. Here, our work reveals an alternative strategy based on subcellular localization, with SNIPE specifically cleaving foreign DNA by localizing to the cell membrane and interacting with proteins associated with phage genome injection (Fig. 6).

**Figure 6:**
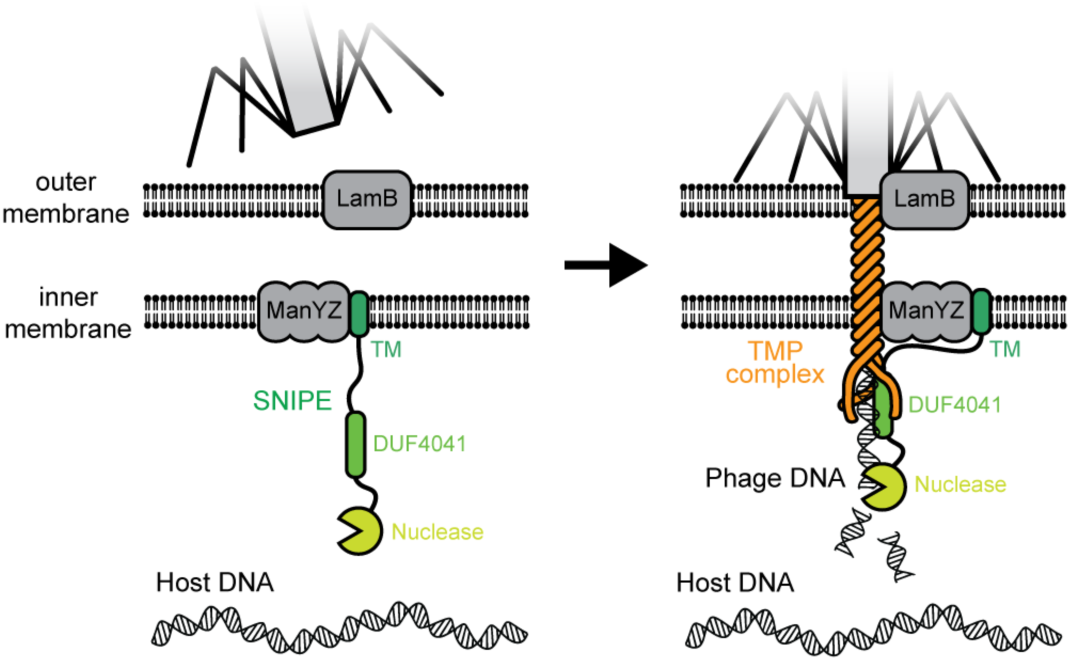
Model for SNIPE-mediated defense against λ. The N-terminal region of SNIPE associates with ManYZ prior to phage infection. This interaction is maintained during phage infection, while the DUF4041 binds to the tape measure protein complex and phage DNA, positioning the nuclease domain to cleave phage DNA. Host DNA is not cleaved by SNIPE during this process. The depiction of the tape measure protein complex is speculative given that it has not been structurally characterized experimentally or *in silico*.

Our proximity labeling studies indicated that SNIPE associates with the mannose permease complex, which is used by λ to promote genome injection, prior to an infection. Then, during infection, SNIPE also associates with the tape measure protein, which has been implicated in forming the putative channel that is required for λ DNA to enter the cell^42,43^. The relative contributions of these two protein complexes to SNIPE defense is not yet clear. However, our finding that SNIPE can defend against a range of siphoviruses independent of ManYZ, along with the identification of SNIPE mutations that enhanced defense against Bas14 in a manner dependent on the sequence of its tape measure protein, indicates that the putative interaction with the tape measure protein may be most critical. Future structural studies are needed to elucidate the detailed molecular interactions that underlie SNIPE-based defense. Such studies also promise to reveal how SNIPE’s nuclease domain is shielded from the host chromosome, which may contact the membrane at some locations^44^, and positioned to only degrade phage DNA as it transits into the cell.

SNIPE homologs are extremely broadly distributed across the bacterial kingdom (Fig. 5A). Notably, although the N-terminal regions of these homologs are not conserved, nearly two-thirds feature a predicted TM domain and the other third include sequences known to target proteins to membranes, such as pleckstrin homology domains. Thus, we anticipate that the SNIPE homologs will, in general, localize to cell membranes and also degrade phage DNA as it is injected using the highly conserved nuclease domains.

Although most anti-phage defense systems act exclusively in the cytoplasm, there are now a handful of membrane-localized systems that target phage DNA. One example is the Zorya system, which likely localizes to phage genome injection sites by sensing perturbations to the peptidoglycan layer and inner membrane, though the exact mechanism underlying its phage detection and specificity remains unclear^45^. Zorya harbors a nuclease domain that may directly cleave phage DNA during genome injection, but whether Zorya exclusively targets phage DNA is complicated by studies arguing that Zorya acts via abortive infection rather than direct defense^37,46^. The Kiwa defense system also localizes to the cell membrane where it is proposed to recognize membrane perturbations associated with phage genome injection, leading to a DNA-binding activity that disrupts phage DNA replication^47,48^. There are also several super-infection exclusion mechanisms, i.e., mechanisms used by a phage to prevent immediate subsequent infection by the same phage, that act at the cell membrane. For instance, the membrane protein gp15 encoded by phage HK97 prevents genome injection in a manner that depends on tape measure protein and certain proteins in the periplasm and inner membrane, though the mechanism of inhibition remains unknown^49,50^.

The importance of systems that block viral entry into the cytoplasm is also exemplified by interferon-induced transmembrane proteins (IFITMs) in eukaryotes^51^. Although the mechanisms of many IFITMs remain unclear, studies suggest that human IFITM3 alters membrane stiffness by enhancing cholesterol production and lipid sorting, trapping viruses in a state of incomplete membrane fusion when they are attempting to enter the host cytoplasm^52–54^. By targeting this conserved aspect of viral entry, IFITM3 helps inhibit infection by a wide range of eukaryotic viruses, including influenza and SARS-CoV^55,56^. Taken together, IFITMs and defense systems like SNIPE underscore the critical role of innate immune mechanisms that block viruses at the early step of genome transfer into host cells.

## Methods

### Growth conditions and strains

All strains, phages, plasmids, and oligos used in this study are listed in Table S1. Sequences of all plasmids were confirmed by whole plasmid sequencing. All bacterial strains were *E. coli* MG1655 derivatives unless noted otherwise. All λ strains in this study had mutations in the cI repressor and/or operator. Phages refered to as simply "λ" correspond to λ_vir_, which cannot establish lysogeny due to operator mutations and a frameshift in *cI*. The generalist λ mutant was derived from λ *cI*26, which has a frameshift in *cI* and is therefore incapable of establishing lysogeny. λ*^parS^* harbors the *cI*857 mutation, which produces a temperature-sensitive repressor that is functional at 30 °C but not at 37 °C.

All bacterial strains were grown in LB or M9 medium (6.4 g L^-1^ Na_2_HPO_4_-7H_2_O, 1.5 g L^-1^ KH_2_PO_4_, 0.25 g L^-1^ NaCl, 0.5 g L^-1^ NH_4_Cl, 0.1% casamino acids, 25 mM MgSO_4_, and 0.1 mM CaCl_2_) at 37 °C unless specified otherwise. Antibiotics were used at the following concentrations (liquid, plates): carbenicillin (carb) (50, 100 μg mL^-1^), chloramphenicol (chlor) (20, 30 μg mL^-1^), kanamycin (kan) (30, 50 µg mL^-1^).

Phages were amplified by back-diluting overnight bacterial cultures to ∼0.05 OD_600_ in 1 mL LB and inoculating with a liquid phage stock at ∼0.1 MOI or a single phage plaque. Phage amplification was performed at 37 °C for 6-20 hours. After this incubation, bacteria were pelleted by centrifugation and the supernatant was transferred to a new tube. This phage supernatant was either vortexed with an additional 100 µL of chloroform or passed through a 0.2 µm filter to remove remaining bacteria.

MacConkey agar plates were prepared using MacConkey agar powder (Difco) and 1% w/v mannose sugar. Strains were streaked out on these plates and grown at 30 °C for 12 hours prior to imaging.

## SNIPE plasmids

SNIPE-GFP was generated by amplifying sfGFP-C1 (Addgene #54579) with oDS6 and oDS7 and pKVS45-PD-λ-1 with oDS5 and oDS8 followed by Gibson assembly. P_van_-SNIPE was generated by amplifying pKVS45-PD-λ-1 with oDS30 and oDS31 and the P_van_ vector followed by Gibson assembly. pBAD*-*SNIPE-mCherry, the parent plasmid for pBAD-SNIPE-mCherry with ΔTM, ΔDUF4041, and/or E414A mutations, was generated by amplifying pKVS45-PD-λ-1 with oDS473 and oDS499, mCherry-pBAD (Addgene #54630) with oDS472 to oDS500, and a pBAD vector with oDS498 and oDS501, followed by Gibson assembly. SNIPE-TurboID-FLAG was generated by amplifying the TurboID-His6_pET21a plasmid (Addgene #107177) using oDS665 and oDS668 and amplifying SNIPE-FLAG with oDS666 and oDS667 followed by Gibson assembly. SNIPE-FLAG was generated by amplifying pKVS45-PD-λ-1 with oDS3 and oDS4, followed by Gibson assembly.

To generate SNIPE(ΔTM) plasmids, parent plasmids were amplified with oDS47 and oDS48 followed by Gibson assembly. Similarly, SNIPE(ΔDUF4041) plasmids were generated by amplifying parent plasmids with oDS51 and oDS52 followed by Gibson assembly. The E414A point mutation was introduced by amplifying parent plasmids with oDS92 and oDS93, followed by Gibson assembly. Other point mutations were generated by amplifying pKVS45-PD-λ-1 with the following primers: oDS923 and oDS924 (L253H); oDS925 and oDS926 (E264K); oDS927 and oDS928 (K305N); oDS929 and oDS930 (E223K); oDS931 and oDS932 (T319A); oDS933 and oDS934 (W257R); oDS935 and oDS936 (I308V); and oDS1075 and oDS1076 (N250amber/pBPA). These PCR products were then circularized by ligation with T4 DNA ligase.

Plasmids with different combinations of the DUF4041, linker, and/or catalytically inactive GIY-YIG nuclease domain fused to GFP were generated by amplifying SNIPE-GFP(E414A) with the following primers: oDS138 and oDS139 (DUF4041-Linker-Nuclease); oDS140 and oDS141 (Linker-Nuclease); and oDS142 and oDS143 (Nuclease). These PCR products were then circularized by Gibson assembly. Two additional combinations were made by amplifying the DUF4041-Linker-Nuclease plasmid made above with the following primers: oDS144 and oDS145 (DUF4041); and oDS146 and 147 (DUF4041-Linker). These PCR products were then circularized by Gibson assembly.

A PhoA-LacZα fusion protein was inserted into SNIPE(E414A) at different locations as described previously^19^. Specifically, the PhoA-LacZα fusion was amplified from pKTop::tse5-CT (Addgene #192955) with the following primers for insertion at different amino acids of SNIPE: oDS412 and oDS413 (SNIPE a.a. 1); oDS416 and oDS417 (SNIPE a.a. 30); oDS420 and oDS421 (SNIPE a.a. 120); and oDS424 and oDS425 (SNIPE a.a. 500). Additionally, P_van_-SNIPE(E414A) was amplified with the following primers: oDS411 and oDS414 (SNIPE a.a. 1); oDS415 and oDS418 (SNIPE a.a. 30); oDS419 and oDS422 (SNIPE a.a. 120); and oDS423 and oDS426 (SNIPE a.a. 500). The two PCR products for a specific insertion point were then assembled with Gibson assembly.

## Other plamids

P_van_-manXYZ was generated by amplifying the *manXYZ* operon from *E. coli* MG1655 with oDS171 and oDS172, and amplifying the P_van_ vector with oDS170 and oDS173, followed by Gibson assembly. Next, P_van_-manXYZ(TurboID-manZ) was generated by amplifying TurboID from TurboID-His6_pET21a with oDS810 and oDS811, and amplifying P_van_-manXYZ with oDS809 and oDS812, followed by Gibson assembly.

To generate P_van_-ftsH-TurboID, the P_van_ vector was amplified with oDS883 and oDS891, ftsH was amplified from *E. coli* MG1655 with oDS892 and oDS893, and TurboID was amplified from TurboID-His6_pET21a with oDS882 and oDS887, followed by Gibson assembly. To generate P_van_-proW-TurboID, the P_van_ vector was amplified with oDS883 and oDS897, proW was amplified from *E. coli* MG1655 with oDS898 and oDS899, and TurboID was amplified from TurboID-His6_pET21a with oDS882 and oDS887, followed by Gibson assembly.

To generate pBAD-EcoRI R-mCherry, EcoRI R was amplified from a the pOpen-EcoRI R plasmid (Addgene #165504) with oDS505 and oDS908, mCherry was amplified from pBAD-SNIPE-mCherry with oDS909 and oDS910, and the pBAD vector was amplified with oDS504 and oDS911; these products were then Gibson assembled.

To generate P_van_-malF(TM1-2)-GFP-fis, fis was amplified from *E. coli* MG1655 with oDS1115 and oDS1116, malF(TM1-2) was amplified from *E. coli* MG1655 with oDS149 and oDS1112, GFP was amplified from sfGFP-C1 (Addgene #54579) with oDS1113 and oDS1114, and the P_van_ backbone was amplified with oDS148 and oDS1117, followed by Gibson assembly.

### *E. coli* gene deletions

*ΔmanYZ*, *ΔmanXYZ*, and *ΔlamB* strains were generated using the lambda red recombination protocol derived from Datsenko and Wanner^57^. Briefly, pKD4 was amplified with oDS94 and oDS95 to generate a Δ*manYZ*::kan^R^ product, oDS95 and oDS174 to generate a Δ*manXYZ*::kan^R^ product, and oDS96 and oDS97 to generate a Δ*lamB*::kan^R^ product. These products were electroporated into either *E. coli* MG1655 or ECOR13 that contained a plasmid that can express lambda red proteins (pKD46). Kan^R^ colonies that contained putative recombination events between the kan^R^ cassette and the gene or operon of interest were isolated and confirmed with junction PCR.

## Phage and bacterial spotting assays

To prepare plates for phage spotting assays, 30 µL of a bacterial overnight culture was added to 4 mL of molten LB + 0.5% agar, and this mixture was then poured onto LB + 1.2% agar plates.

After the top agar solidified, ten-fold serial dilutions of a phage stock were made and 2 µL of each dilution were pipetted onto the plate with a multichannel pipette. Plates were incubated at 37 °C for 8-20 hours prior to imaging. Images are representative of at least two independent biological replicates. EOP values were assessed qualitatively given that strong defense prevents formation of individual plaques. For one exception, Extended Data Fig. 7b, we used the top agar overlay method with different strains of interest and quantified plaques for three independent biological replicates.

To perform bacterial spotting assays, overnight cultures of bacteria were back-diluted to 1 OD_600_, ten-fold serial dilutions of this bacterial stock were made, and 3 µL of each dilution were spotted onto LB + 1.2% agar + 1, 40, or 80 µM vanillate plates. Plates were incubated at 37 °C for ∼12 hours prior to imaging. Images are representative of three independent biological replicates.

## Growth curve assays

Overnight cultures were back-diluted to 0.01 OD_600_ and grown to mid-log phase in LB at 37°C, then back-diluted to 0.1 OD_600_. 100 µL aliquots of these cell solutions were added to wells of a 96-well plate. 10 µL of different phage dilutions were then added to generate different MOIs.

Samples were incubated at 37 °C with orbital shaking on a plate reader (Biotek) for 8 hours, with OD_600_ measurements every 10 minutes for 10 hours.

## Microscopy

To prepare cells for microscopy, overnight cultures were back-diluted to 0.05 OD_600_ and grown in LB at 37 °C until cells reached mid-exponential phase (0.3 to 0.4 OD_600_) unless specified otherwise.

Cells were then concentrated to 1-1.5 OD_600_ to have a high cell density for microscopy. LB + 1.5% Ultrapure Agarose (Invitrogen 16500-100) was melted and 600 µL was added to a 22mm x 22mm No. 1.5 coverslip (VWR) and another identical coverslip was immediately placed on top. After the agarose pad solidified, the top coverslip was removed and 0.2 µL of cells were spotted onto the pad. For a given experiment, multiple bacterial strains were spotted onto the same agarose pad at different positions. After the spots soaked into the agarose pad, a 50mm x 22mm No. 1.5 coverslip (VWR) was placed on top of the pad. Samples were imaged with phase contrast and epifluorescence channels on a Zeiss Observer Z1 microscope with a 100X/1.4 oil immersion objective and Colibri illumination system. Samples were kept at 37 °C during imaging by using a XLmulti S incubator (Pecon) and Heating Unit XL and TempModule S (Zeiss). An Orca Flast 4.0 camera (Hamamatsu) and Metamorph (Molecular Devides) were used for imaging. Image analysis was performed with Fiji (ImageJ). All samples within an experiment are shown with similar brightness unless otherwise noted.

For SNIPE-GFP localization experiments, mid-exponential phase cells were incubated with 10 µg mL^-1^ DAPI for ten minutes prior to imaging. These cells were imaged with phase contrast, DAPI, and FITC channels. Images are representative of three independent biological replicates. Pearson’s correlation coefficients were calculated in Fiji (ImageJ) using the Coloc 2 plugin, following thresholding of the Phase channel, conversion to a mask, and segmentation with the watershed algorithm.

For experiments with Gam-GFP, cells were grown to mid-exponential phase in M9 media + 1% glucose (to repress SNIPE-mCherry or EcoRI R-mCherry) + 10 µg mL^-1^ aTc (to express Gam-GFP) + carb + chlor. Cells were then washed with and resuspended in M9 media + 0.25% arabinose (to express SNIPE-mCherry or EcoRI R-mCherry) and incubated at 37 °C for 40 minutes. DAPI was then added to 10 µg mL^-1^ final concentration and cells were incubated at 37°C for another ten minutes. These cells were then imaged on a M9 media + 1.5% agarose pad with FITC, rhodamine, DAPI, and phase constrast channels. Images are representative of three biological replicates. Maximum pixel intensity values were quantified in Fiji (ImageJ) after thresholding the Phase channel, converting it to a binary mask, and segmenting cells using the watershed algorithm. Individual cells were then analyzed with the Analyze Particles → Measure function.

For experiments involving infection of CFP-ParB cells with λ*^parS^*phage, cells were grown to mid-exponential phase in LB + 20 µM IPTG (to induce CFP-ParB expression) + carb + chlor at 37 °C. Cells were concentrated to 1.5 OD_600_ in 100 µL of the same media and placed on ice for five minutes. λ*^parS^* was then added at an MOI of ∼6 and incubated with cells on ice for 30 minutes to facilitate adsorption but prevent genome injection. This phage and cell mixture was then spotted onto a cooled LB + 1.5% agarose pad with 20 µM IPTG, carb, and chlor. This agarose pad was quickly transferred to the microscope, which maintained the samples at 37 °C and thus triggered genome injection. Samples were imaged in phase contrast and CFP channels every 5 minutes for 150 minutes. Quantification of CFP-ParB foci per cell and cell death per infected cell was performed manually. Images are representative of three biological replicates.

For experiments involving induction of λ*^parS^ cI*857 prophages via heat shock, lysogens containing this prophage were grown to mid-exponential phase in LB + 80 µM IPTG (to induce CFP-ParB) + carb + chlor at 30°C to prevent prophage induction. This cell solution was transferred to a 42 °C heat block for five minutes to initiate prophage induction. Cells were then spotted onto a pre-warmed LB + 1.5% agarose pad with 80 µM IPTG, carb, and chlor. This pad was transferred to the microscope, where it was maintained at 37 °C and imaged in phase contrast and CFP channels every 5 minutes for 150 minutes. Remaining cells that were not spotted onto the agarose pad were incubated at 37 °C for 4 hours after which plaque forming units were quantified via spotting assays. Comparison of plaque forming units per OD_600_ of cells that were initially induced yielded values shown in Fig. 2f. Images and phage spotting assays are representative of three independent biological replicates.

To test for direct defense and abortive infection phenotypes, cells were grown to mid-exponential phase in LB + chlor followed by addition of λ at 2 MOI, which ultimately yielded approximately one phage infection event for every two cells. This phage and cell mixture was incubated at 37°C for eight minutes to allow phage adsorption, followed by two rapid washes in LB + chlor to remove unadsorbed phage. This phage and cell mixture was then spotted onto an LB + 1.5% agarose pad with chlor and imaged in phase contrast for 3 hours at 10-minute time intervals. The imaging started exactly 20 minutes after the phage and cells were first mixed. Images and movies are representative of three independent biological replicates.

For experiments involving MalF(TM1-2)-GFP-Fis, cells were grown to mid-log phase in LB and 50 µM vanillate was added. This solution was incubated at 37°C for 30 minutes. DAPI was then added to 10 µg mL^-1^ final concentration and cells were incubated at 37 °C for another ten minutes. These cells were then imaged on a LB media + 1.5% agarose pad with FITC, DAPI, and phase constrast channels. Images are representative of three independent biological replicates.

## PhoA-LacZ**α** topology assay

Membrane topology assays were performed as previously described^19,58^, using the *E. coli* DH5α strain, which is naturally *ΔphoA* and is capable of α-galactosidase α-complementation (provided by LacZα). Specifically, three independent overnight cultures per strain were back diluted to 0.05 OD_600_ in 3 mL LB + 200 µM vanillate to induce P_van_-SNIPE constructs and grown to late log phase (∼0.8 OD_600_). To test PhoA activity, 1 mL of each sample was washed once in 1 mL P buffer (1 M Tris-HCl pH 8.0 and 0.1 mM ZnCl_2_) and resuspended in 1 mL P buffer. 50 µL of chloroform and 50 µL of 0.1% SDS was added to each sample and vortexed for 5 seconds to permeabilize cells. This solution was incubated at 30 °C while chloroform separated from the aqueous layer. 150 µL of the aqueous layer was transferred to a 96-well plate. The enzymatic reaction was initiated by adding 18 µL of p-nitrophenylphosphate solution (Sigma N7653) to each sample. Samples were incubated at 30 °C with orbital shaking on a plate reader (Biotek) for 2 hours, with OD_405_ measurements every 2 minutes. OD_600_ was also measured to calculate the number of cells in each sample.

In parallel, to test LacZ activity, 1 mL of each late-exponential phase cultures described above was washed once with 1 mL of Z buffer (60 mM Na_2_HPO_4_, 40 mM NaH_2_PO_4_, 10 mM KCl, 1 mM MgSO_4_, 50 mM α-mercaptoethanol added on day of experiment) and resuspended in 1 mL Z buffer. 50 µL of chloroform and 50 µL of 0.1% SDS was added to each sample and vortexed for 5 seconds to permeabilize cells. This solution was incubated at 30 °C while chloroform separated from the aqueous layer. 150 µL of the aqueous layer was transferred to a 96-well plate. The enzymatic reaction was initiated by adding 18 µL of 4 mg/mL o-nitrophenyl galactopyranoside (Sigma N1127) to each sample. Samples were incubated at 30 °C with orbital shaking on a plate reader (Biotek) for 2 hours, with OD_420_ measurements every 2 minutes. OD_600_ was also measured to calculate the number of cells in each sample.

To measure the enzymatic activities of PhoA and LacZ, the difference in OD_405_ or OD_420_ measurements between 0 and 60 minutes was calculated and calibrated to the starting OD_600_ of the sample to provide enzymatic activity per cell. These analyses were performed separately for the three independent biological replicates prior to calculating averages and standard deviations.

## Membrane fractionation and immunoblots

Overnight bacterial cultures were back-diluted in LB and grown to 0.6 OD_600_ at 37°C. Cells were pelleted by centrifugation and resuspended in Buffer A (150 mM NaCl, 50 mM HEPES pH 7) with EDTA-free protease inhibitor (Roche), 5 µL (150 kU) Ready-Lyse (Biosearch technologies), 5 µL (125 U) benzonase (Millipore), and 2 mM MgCl_2_. For measurements of protein levels in lysates, this cell solution was lysed in 1x Laemmli buffer with gentle heating at 50 °C, followed by electroporation and immunoblots as described below. For membrane fractionation, the cell solution was incubated on ice for 30 minutes to facilitate degradation of the peptidoglycan layer and then lysed by sonication. Lysate was centrifuged at 10,000 *g* to pellet debris, and the supernatant was transfered to a new tube and centrifuged at 140,000 *g* to pellet membranes. The supernatant from this spin was saved as the cytoplasmic fraction. The membrane pellet was briefly rinsed with Buffer A and resuspended in 2 mL of Buffer A with a Dounce homogenizer.

Samples were normalized by protein concentration, mixed with 4x Laemmli buffer, and centrifuged at 13,000 *g* for ten minutes to pellet any debris. 25 µL of each sample was subjected to electrophoresis with a 4-20% polyacrylamide gel and transferred to a 0.2 µm polyvinylidene difluoride membrane. Anti-GFP (Invitrogen A11120), anti-DnaK (AssayPro 32857-05111), and anti-OmpC (Bioss Antibodies bs20213R) antibodies were used at a final concentration of 1:1000 and SuperSignal West Femto Maximum Sensitivity Substrate (ThermoFisher) was used to develop blots. Blots were imaged using a ChemiDoc Imaging system (Bio-Rad). Images shown are representative of two biological replicates.

## Adsorption assay

Overnight bacterial cultures were back-diluted to 0.05 OD_600_ in 1 mL of LB and grown to 0.5 OD_600_ at 37 °C. These cultures were infected with λ at an MOI of 0.1 and incubated at 37 °C for 15 minutes. Samples were then centrifuged at 10,000 *g* for three minutes and the phage-containing supernatant was transferred to a new tube containing 100 µL chloroform, which was then vortexed. PFU per µL were calculated using the top agar overlay method with *E. coli* MG1655. Percent adsorption was determined relative to a simultaneous control sample that contained growth medium but no cells. Data represent the averages and standard deviations of three independent biological replicates.

## Injection of radiolabeled phage DNA

A protocol to generate λ with ^32^P-labeled DNA was designed using information from Hershey & Chase 1952 and Stent & Fuerst 1955^23,59^. First, low-phosphate H-media (20 mM KCl, 85 mM NaCl, 20 mM NH4Cl, 1 mM MgSO4, 50 µM CaCl2, 70 mM sodium lactate, 0.2% glycerol, 0.05% bacto-peptone (Difco), and 0.05% bacto-casamino acids (Difco)) was prepared as a 2x concentrate and diluted in water and 4% agar to generate molten H media + 1.2% agar and molten H media + 0.5% agar. 5 mL of H media + 1.2% agar was poured into a 60 mm petri dish and allowed to solidify. Next, 3 mL of the molten H media + 0.5% agar was mixed with 15 µL of *Δdam E. coli* overnight culture (to prevent Dam methylation of DNA in this strain, the purpose of which is described below), 10,000 PFU λ, and 300 µCi ^32^P (Revity). This mixture was immediately plated onto the H media 1.5% agar plate, which was incubated at 37 °C overnight.

This protocol produced confluent lysis of the *E. coli*, which permitted recovery of much higher phage titers than similar attempts at phage amplification in liquid H media.

To recover ^32^P-labeled phage, 1 mL of FM buffer (20 mM Tris-HCl pH 7.4, 100 mM NaCl, 10 mM MgSO4) was gently added to the top of the plate and incubated for 3 hours at room temperature. This solution was then aspirated off of the plate and transferred to a microfuge tube. RNaseA was added at 10 µg mL^-1^ and incubated for five minutes at room temperature to degrade ^32^P-labeled RNA. This solution was centrifuged at 10,000 *g* to remove bacterial debris, transferred to a new tube with 10% volume chloroform, vortexed, and centrifuged again. The supernatant was transferred to a 0.5 mL 50 kDa MWCO spin concentrator (Amicon) and repeatedly spin concentrated and resuspended in FM buffer to quickly remove excess ^32^P present in the media. This protocol typically yielded 200 µL of 10^6^ PFU µL^-1^ λ.

To infect cells with ^32^P-labeled λ, overnight cultures of different *E. coli* strains (which contained functional Dam and thus methylate their DNA) were back-diluted to 0.1 OD_600_ in 1 mL of LB and grown at 37°C until they reached 0.4 OD_600_. These cell solutions were then centrifuged, resuspended in 100 µL of LB, and placed on ice for five minutes. Next, the entire volume of radiolabeled phage was equally distributed across the bacterial samples, which yielded an MOI of approximately 0.1. These phage and cell mixtures were incubated on ice for 20 minutes to facilitate adsorption, then 37 °C for 15 minutes to trigger genome injection. Cells were then spun down and resuspended in 1 mL of ice-cold PBS, and this wash step was repeated a second time, effectively removing unadsorbed phage particles. Cells were then spun down and resuspended in 25 µL ice-cold 1x rCutSmart buffer + 0.5 µL (15 kU) Ready-Lyse (Biosearch Technologies) and incubated on ice for 5 minutes. This solution was flash-frozen with liquid nitrogen and thawed three times to lyse cells. 0.5 µL (10 U) of DpnI (NEB) was added to each sample and incubated at 37 °C for five minutes to degrade the Dam-methylated *E. coli* DNA and thus decrease the viscosity of the sample (notably, because phage were amplified in a *Δdam* strain, they did not have methylated DNA and phage DNA was thus resistant to DpnI-mediated degradation).

Two controls were prepared at this time. First, one aliquot of empty vector cells that had been infected with ^32^P-labeled λ was treated with 1 µL (25 U) of benzonase (Millipore) and incubated at 37 °C for five minutes to degrade all DNA in that sample. Additionally, a second sample of empty vector cells that had been infected with ^32^P-labeled λ was mixed with a sample of SNIPE-expressing cell lysate that had not been infected with phage, and this mixture was incubated at 37°C for five minutes. This effectively tested if SNIPE is capable of cleaving phage DNA in cell lysates (i.e., after genome injection had already occurred), which did not turn out to be the case.

To measure the length distributions of ^32^P-labeled DNA in these samples, these lysates were mixed with 4x Laemmli buffer and centrifuged at 13,000 *g* for ten minutes to pellet any remaining bacterial debris. 25 µL of each sample was loaded into a 4-20% pre-cast polyacrylamide gel (Bio-Rad). This gel was run at 100V for 20 minutes and then 150 V for 25 minutes. The gel was incubated with a phorphorscreen and imaged using a Typhoon imager (GE Healthcare). Data for two independent biological replicates are shown.

## Proximity labeling with TurboID

Proximity labeling assays were performed as previously described^27^, with some modifications. Overnight bacterial cultures were back diluted to 0.1 OD_600_ in 50 mL of LB + appropriate antibiotics +/-vanillate and grown at 37 °C to 0.4 OD_600_. The P_van_ promoter was used to express various constructs (TurboID-ManZ, FtsH-TurboID, and ProW-TurboID). Leaky expression under this promoter in the absence of vanillate was sufficient to express TurboID-ManZ, but not FtsH-TurboID, or ProW-TurboID. Therefore, 0 µM vanillate was used for some experiments (Extended Data Fig. 4b, f, g, i) and 5 µM vanillate was used for others (Fig. 3c, Extended Data Fig. 4c-e).

Following cell growth, these samples were then centrifuged, and cells were resuspended in 1.5 mL putrescine buffer (10 mM Tris-HCl pH 7.4, 10 µM MgCl2, 10 mM putrescine dihydrochloride). Previous studies demonstrated that polyamines such as putrescine "lock" λ DNA in the phage capsid, such that the genome injection apparatus is probably able to assemble but DNA injection does not complete^60,61^; this phenomenon helped stall otherwise transient genome injection events to permit better proximity labeling of phage proteins. Biotin (dissolved in DMSO to 100 mM) was added at 500 µM final concentration, λ was added at an MOI of 40, and these samples were placed at 37 °C for fifteen minutes to facilitate biotinylation of phage and host proteins.

After this incubation step, samples were washed three times by centrifugation and resuspension of cells in 1 mL ice-cold putrescine buffer + 1 µL (25 U) benzonase (Millipore). Finally, cells were pelleted and resuspended in 1.5 mL RIPA buffer (50 mM Tris-HCl pH 7.4, 150 mM NaCl, 1% Triton X-100, 0.5% sodium deoxycholate, 0.1% SDS) + EDTA-free protease inhibitor (Roche) + 1 µL (25 U) benzonase (Millipore) + 1 µL (30 kU) Ready-Lyse (Biosearch Technologies). Cells were lysed via sonication and then spun at 15,000 *g* for ten minutes to pellet cell debris. The supernatant was transferred to a fresh tube with 75 µL of streptavidin magnetic beads (Invitrogen 65002) that had been pre-equilibrated in RIPA buffer. This mixture was incubated on an end-over-end rotor at 4 °C overnight.

Streptavidin beads were subsequently washed twice with each of the following buffers: RIPA buffer, 1 M KCl, 100 mM Na2CO3, Urea buffer (2 M urea, 10 mM Tris-HCl pH 7.4), and PBS. On-bead reduction, trypsin digestion, and LC–MS/MS were done by the MIT Biopolymers and Proteomics Core. Specifically, proteins were reduced with 10 mM dithiothreitol (Sigma) for 1 hour at 56 °C, followed by alkylation with 20 mM iodoacetamide (Sigma) for 1 hour at 25 °C in the dark. Digestion was performed overnight using modified trypsin (Promega) in 100 mM ammonium bicarbonate (pH 8) at an enzyme-to-substrate ratio of 1:50. The digestion was terminated by the addition of formic acid (99.9%, Sigma). The resulting peptides were desalted using Pierce Peptide Desalting Spin Columns (Thermo) and subsequently lyophilized. Peptide separation was carried out on a PepMap RSLC C18 column (Thermo) over a 90-minute gradient using reverse-phase high-performance liquid chromatography (Thermo Ultimate 3000), followed by nano-electrospray ionization and analysis on an Orbitrap Exploris 480 mass spectrometer (Thermo). The mass spectrometer operated in data-dependent acquisition mode, with full-scan parameters set to a resolution of 120,000 across an m/z range of 375–1600 and a maximum ion injection time of 25 ms. MS/MS acquisition was performed for as many precursor ions as possible within a two-second cycle, using a resolution of 30,000, a normalized collision energy (NCE) of 28, and a dynamic exclusion window of 20 seconds. Peptides were mapped to proteins from *E. coli* MG1655, proteins from λ, and proteins of interest such as SNIPE-GFP and TurboID-ManZ. To compute ratios between spectral counts from different samples, a psuedocount was first added to each spectral count value. Additionally, ratios were only computed for proteins in which the number of spectral counts was ≥50 for at least one of the samples.

## Error-prone PCR and SNIPE mutant selection

SNIPE was mutagenized using error-prone PCR-based mutagenesis, as previously described^62^. Different regions of SNIPE (corresponding to amino acids 1-190, 191-330, and 331-500) were amplified using Taq polymerase (NEB) and 0.5 mM MnCl_2_ was added to the reaction as the mutagenic agent. PCR products were treated with DpnI, gel extracted, and separately cloned into the original pKVS45-PD-λ-1 plasmid using Gibson assembly with the goal of creating three pools with mutations in these different regions. Gibson products were column purified and electroporated into *E. coli* MG1655. ∼10 million independent transformants were recovered for each pool, which were then combined to generate a mutagenized library of ∼30 million SNIPE mutants.

To select SNIPE mutants with enhanced defense against Bas14, the SNIPE mutant library was grown to saturation overnight, 50 µL of this culture was mixed with Bas14 at an MOI of 1, and this solution was immediately plated on LB + 1.5% agar + chlor plates. After incubation overnight at 37 °C, Bas14-resistant colonies were picked, streaked for single colonies, grown into cultures, miniprepped, and the resulting plasmids were transformed into *E. coli* MG1655. Plasmids that conferred protection against Bas14 were sequenced to identify SNIPE mutations.

## Isolation and sequencing of phage escapers

Bas14 escapers were selected by mixing 30 µL of *SNIPE(L253H E264K K305N)* overnight culture and 10^9^ PFU Bas14 with molten LB + 0.5% agar and using the top agar overlay method. Individual plaques were amplified in the *SNIPE(L253H E264K K305N)* strain and single plaques were isolated and amplified again to generate isogenic phage stocks.

Phage DNA was extracted by treating 200 µL of phage stock (over 10^7^ PFU µL^-1^) with 0.2 U DNaseI and 0.05 mg mL^-1^ RNaseA at 37°C for 30 minutes. 10 mM EDTA was then added to inactive these nucleases. This solution was then incubated with Proteinase K at 50 °C for 30 minutes to disrupt phage particles and release DNA, which was then isolated by ethanol precipitation.

To prepare Illumina sequencing libraries, 200 ng of phage DNA was sheared in a Diagenode Bioruptor 300 sonicator water bath for 20 x 30 second cycles at maximum intensity. Sheared gDNA was prepared for Illumina sequencing with the NEBNext Ultra II DNA library preparation kit and sequenced on an Illumina MiSeq at the MIT BioMicroCenter. Illumina reads were assembled to the Bas14 reference genome using Geneious Prime 2025.3.

## Tn-Seq

Tn-Seq of cells in the presence or absence of phage infection was performed as previously described^63^. A pool of cells haboring random transposon insertions^64^, each with a unique barcode and a kan^R^ cassette, was grown to late-log phase and aliquots containing ∼10^7^ cells were prepared. Different phages were added to separate tubes at ∼1 MOI, and LB was added to a no phage control. These samples were then immediately plated on LB + kan plates and grown overnight at 37 °C. Colonies were scrapped off of the plates and gDNA from 1 OD_600_ unit of cells per sample was extracted with a DNeasy Blood and Tissue Kit (Qiagen). 85 ng of DNA from each sample was amplified with BarSeq primers (Table S1) to specifically amplify transposon barcodes with flanking Illumina adapter and index sequences. PCR products were pooled, treated with DpnI, run on an agarose gel, and gel extracted. The pooled library was sequenced on an Illumina MiSeq at the MIT BioMicroCenter.

To count the number of reads for each barcode, we used previously written scripts^64^ available at https://bitbucket.org/berkeleylab/feba. We log_2_-normalized read counts within each condition by calculating for each read count, 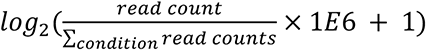. Then, to identify enriched barcodes, for each phage we fit a linear regression between the log_2_-normalized read counts of the no phage condition (independent variable) and the log_2_-normalized read counts of the phage condition (dependent variable). For each barcode, the residual from the linear regression represented its deviation from expectation. We then mapped barcodes to genes using previously described mapping information^64^ and averaged residuals for each gene to get log_2_ enrichment scores.

## Unnatural amino acid crosslinking

Overnight bacterial cultures harboring different SNIPE-FLAG constructs and a plasmid with a suppressor tRNA and specialized tRNA synthetase (pSup-pBpaRS-6TRN)^32^ were back diluted to 0.05 OD_600_ in 2 L of LB + 1.2 mM pBPA and grown at 37 °C to 0.4 OD_600_. Cells were then pelleted by centrifugation and resuspended in 25 mL of putrescine buffer (10 mM Tris-HCl pH 7.4, 10 µM MgCl2, 10 mM putrescine dihydrochloride). Bas14 was then added at 30 MOI and samples were exposed to UV with a Evoluchem 365 nm LED (HepatoChem) for 15 minutes. Cells were then pelleted by centrifugation and resuspended in 25 mL ice-cold Buffer A (50 mM HEPES pH 7.5, 150 mM NaCl) + EDTA-free protease inhibitor (Roche) + 1 µL (25 U) benzonase (Millipore). Cells were lysed with an LM20 Microfluidizer (Microfluidics) at 18,000 psi and cell debris was pelleted at 10,000 *g* for 30 minutes. 20% Anzergent 3-12 was added to the supernatant to a final concentration of 1% to solubilize membrane proteins. This solution was then spun at 100,000 *g* for 1 hour to pellet unsolubilized membranes, and the supernatant was incubated with 75 µL of Pierce™ anti-FLAG magnetic beads (Thermo) overnight at 4°C. Beads were washed three times with Buffer A + 150 mM NaCl + 0.1% Anzergent 3-12, then washed three times with Buffer A + 150 mM NaCl to remove remaining detergent. On-bead reduction, trypsin digestion, and LC–MS/MS were done by the MIT Biopolymers and Proteomics Core as described above. Peptides were mapped to proteins from *E. coli* MG1655 and Bas14. To compute ratios between spectral counts from different samples, a psuedocount of 0.5 was first added to each spectral count value.

## SNIPE homolog analysis

SNIPE homologs analyzed in this study were originally identified by Vassallo et. al^17^. The original set of 1141 homologs was condensed with MmSeqs2 with a clustering threshold of 0.95 and an alignment coverage threshold of 0.9 to remove highly similar homologs. This condensed list of 512 homologs was then aligned and manually curated to remove homologs that were clearly truncated versions of other homologs in the set, as would be expected from misannotated start codons. The resulting list was comprised of 474 SNIPE homologs and used for analyses in this study.

N-terminal regions of SNIPE were extracted and used in a HHblits search for hits in the Pfam database (PFAM), conserved domain database (cdd) from NCBI, and the Protein Data Bank (PDB). Hits with a p-value of <10^-5^ were considered significant and used for further analysis. These hits were sorted into two sets, one set from SNIPE homologs with 1-2 predicted TMs, and the other set from SNIPE homologs that lacked predicted TMs. Because there were usually multiple significant hits for each SNIPE homolog and many homologs had similar hits, we identified the most common hit within a set, calculated the number of SNIPE homologs containing that hit, and then removed all SNIPE homologs that contained one of these hits from further analysis. This process was iterated multiple times to generate a list of common hits for SNIPE homologs lacking TMs, and a separate list of common hits for the SNIPE homologs containing 1-2 TMs, which is shown in Extended Data Fig. 10a.

To search for structural homologs of N-terminal regions of SNIPE homologs that lacked predicted TMs, AlphaFold2 was used to predict structures for this set of SNIPE homologs. Each predicted structure was manually examined for N-terminal regions that harbored globular domains. To identify structural homologs of these domains, FoldSeek was used to scan the PDB100 database (version 20240101).

## Data availability

Sequencing data is available in the Sequence Read Archive under BioProject PRJNA1231458. Summaries of spectral read counts and raw data for MS/MS of biotinylated or crosslinked proteins were deposited under MassIVE and can be accessed under accession MSV000097285. All other data are available in the manuscript or supplementary materials. Source data are provided with this paper.

## Acknowledgements

We thank T. Zhang, C. Vassallo, P. DeWeirdt, and D. Nguyen for comments on the manuscript. We thank J. Meyer for sharing the generalist λ mutant, I. Golding for sharing λ*^parS^* and CFP-ParB strains, the MIT BioMicroCenter and their staff for support with Illumina sequencing, and the MIT Biopolymers and Proteomics Core and its staff for assisting in mass spectrometry. We thank A. Millman for help with generating the unrooted phylogenetic tree. We thank A. Caruso for reagents and advice on pBPA crosslinking. M.T.L. is an Investigator of the Howard Hughes Medical Institute. This work was also supported by a Helen Hay Whitney Fellowship awarded to D.S.S.

## Author contributions

All experiments were done by D.S.S. Experiments were designed and conceived by D.S.S., I.R., and M.T.L. Bioinformatic analysis was done by D.S.S., C.R.D., and P.C.D. Figure design, manuscript writing, and editing done by D.S.S. and M.T.L. Project supervision and funding provided by M.T.L.

---

**Extended Data Figure 1.**
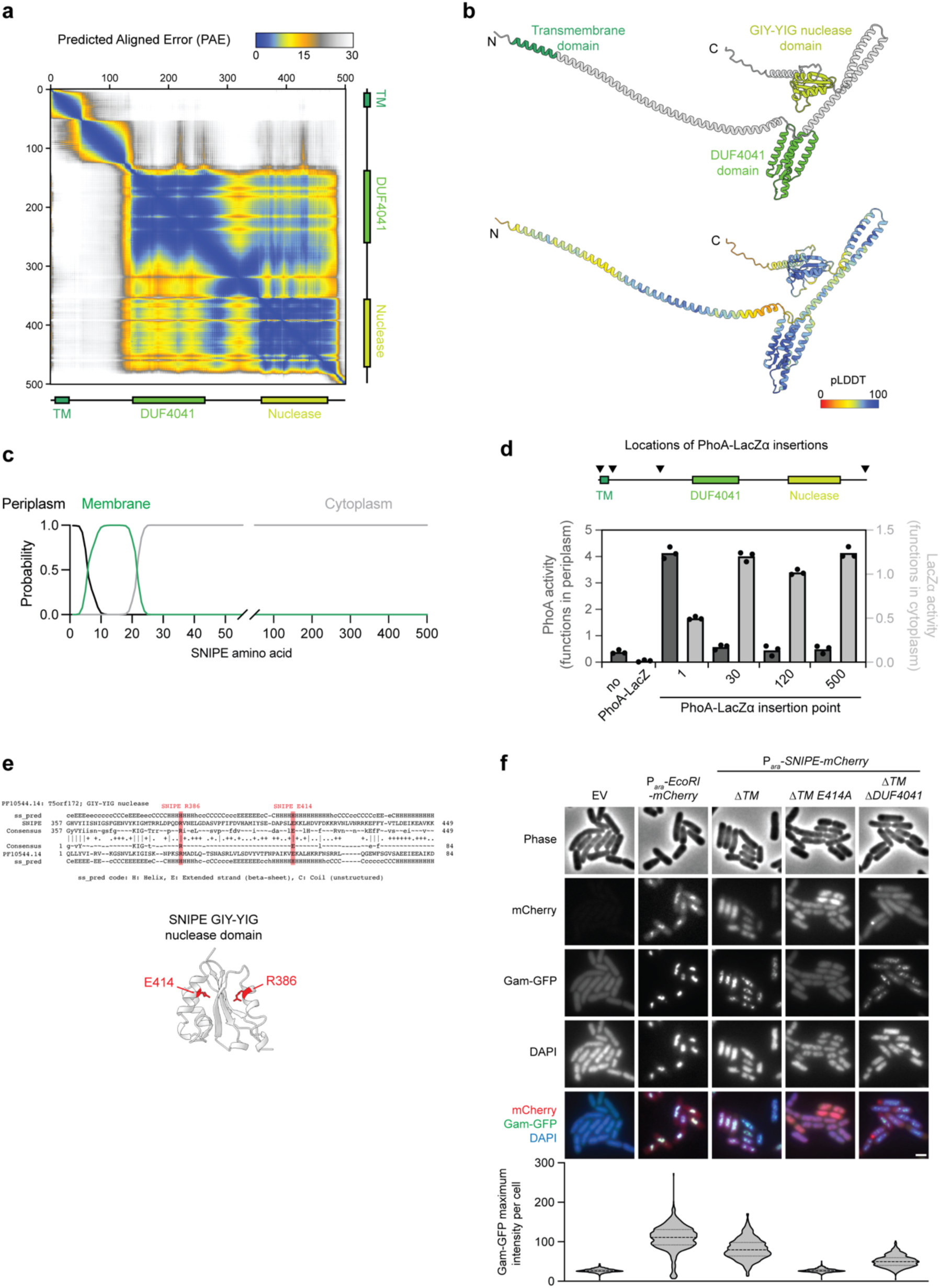
: Predicted structure of SNIPE and membrane topology assay. **(a)** PAE plot for SNIPE structure predicted by AlphaFold3. **(b)** The structure of SNIPE predicted by AlphaFold3, as shown in Fig. 1b, with pLDDT scores shown on the bottom structure. **(c)** Predicted topology of SNIPE residues calculated by DeepTMHMM. **(d)** Insertion points of PhoA-LacZα in SNIPE(E414A) are shown in the schematic. PhoA and LacZ activities were calculated by permeabilizing cells in the presence of colorimetric substrates p-nitrophenylphosphate and o-nitrophenyl galactopyranoside, respectively. Summary of 3 independent replicates. **(e)** HHpred analysis of SNIPE shows that the nuclease domain is a member of the GIY-YIG nuclease family. The structure of the nuclease domain, as predicted by AlphaFold3, is shown on the right. Predicted catalytic residues are highlighted in red. **(f)** Single-frame microscopy of cells expressing an empty vector or P*_ara_*-SNIPE-mCherry constructs, and Gam-GFP, a marker of double strand breaks. The nucleoid was stained with DAPI. A subset of these images is shown in Fig. 1f. Given that Gam-GFP foci often coalesce^20^, the maximum Gam-GFP pixel intensity per cell is shown as a violin plot. Center dashed line, median; top and bottom dashed lines, upper and lower quartiles. n ≥ 431 cells per sample. Scale bar, 1 µm.

**Extended Data Figure 2.**
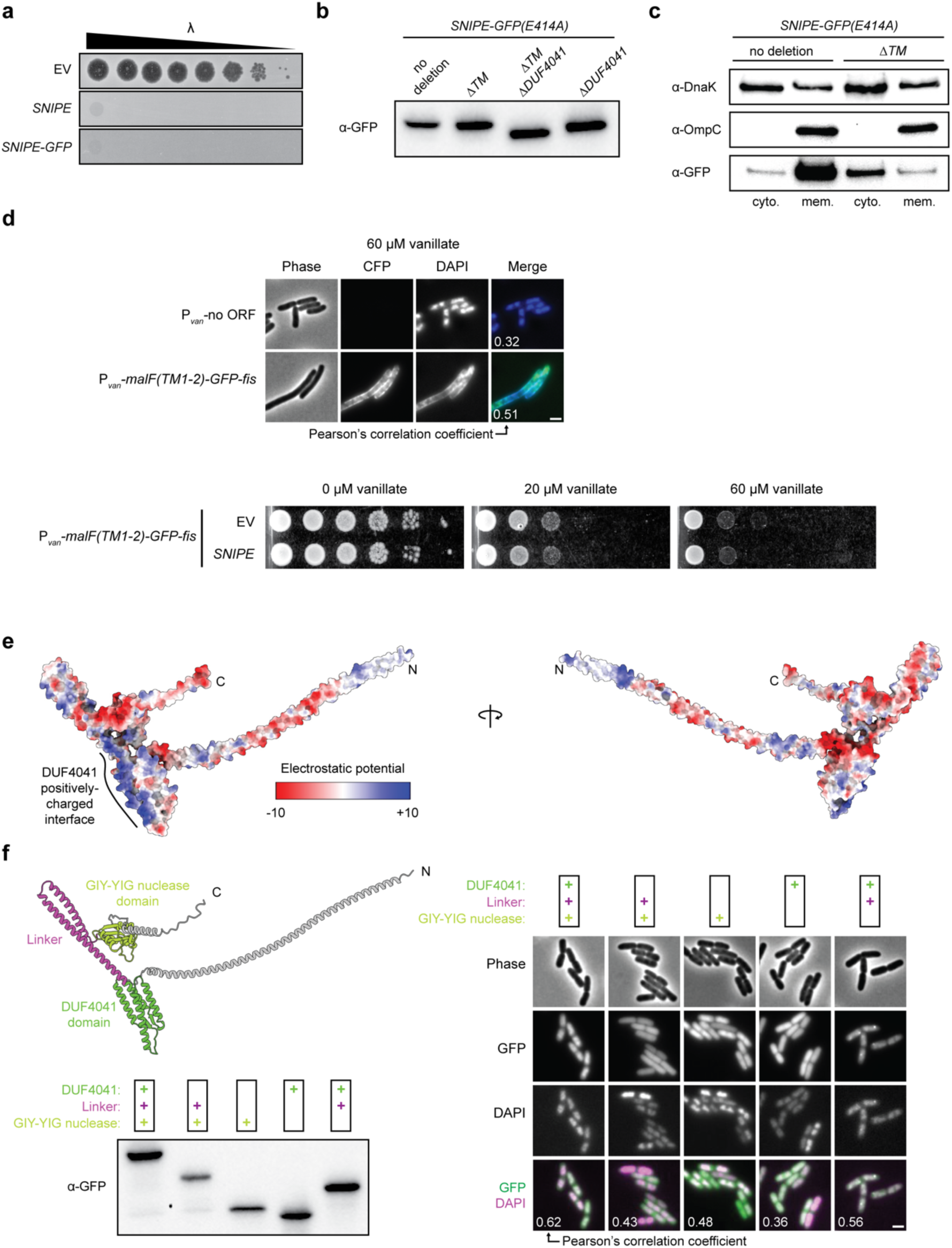
: Localization and nuclease activity of fluorescently tagged SNIPE. **(a)** Serial dilutions of λ were spotted onto lawns of cells expressing an empty vector (EV), SNIPE, or SNIPE-GFP. **(b)** Expression levels of different SNIPE-GFP(E414A) constructs as shown by immunoblots. The same membrane was stained with Coomassie as a loading control. For gel source data, see Supplementary Figure 1a. **(c)** Immunoblots of separated cytoplasm and membrane fractions isolated from cells expressing SNIPE(E414A)-GFP or SNIPE(ΔTM E414A)-GFP. Immunoblots for DnaK, which is mostly cytoplasmic^65^, and OmpC, which is a membrane protein^66^, are provided as controls. The anti-GFP membrane was stained with Coomassie as a loading control. For gel source data, see Supplementary Figure 1b. **(d)** Strains expressing MalF(TM1-2)-GFP-Fis or a negative control plasmid were imaged with fluorescence and phase microscopy. The nucleoid is stained with DAPI. Scale bar, 2 µm. Additionally, toxicity of strains expressing MalF(TM1-2)-GFP-Fis and harboring SNIPE or an empty vector control was assessed with bacterial spotting assays on plates with 0, 20, or 60 µM vanillate. **(e)** Electrostatic surfaces of the AlphaFold3-predicted SNIPE structure. The positively charged surface of the DUF4041 is highlighted. **(f)** The structure of SNIPE predicted by AlphaFold3, with color-coded domains of interest. Different combinations of these domains were fused to GFP and imaged with fluorescence and phase microscopy. The nucleoid was stained with DAPI. All constructs that contained the nuclease domain harbored the catalytically inactive E414A mutation. Scale bar, 2 µm. Expression levels of these constructs were assessed with immunoblots. The same membrane was stained with Coomassie as a loading control. For gel source data, see Supplementary Figure 1a.

**Extended Data Figure 3.**
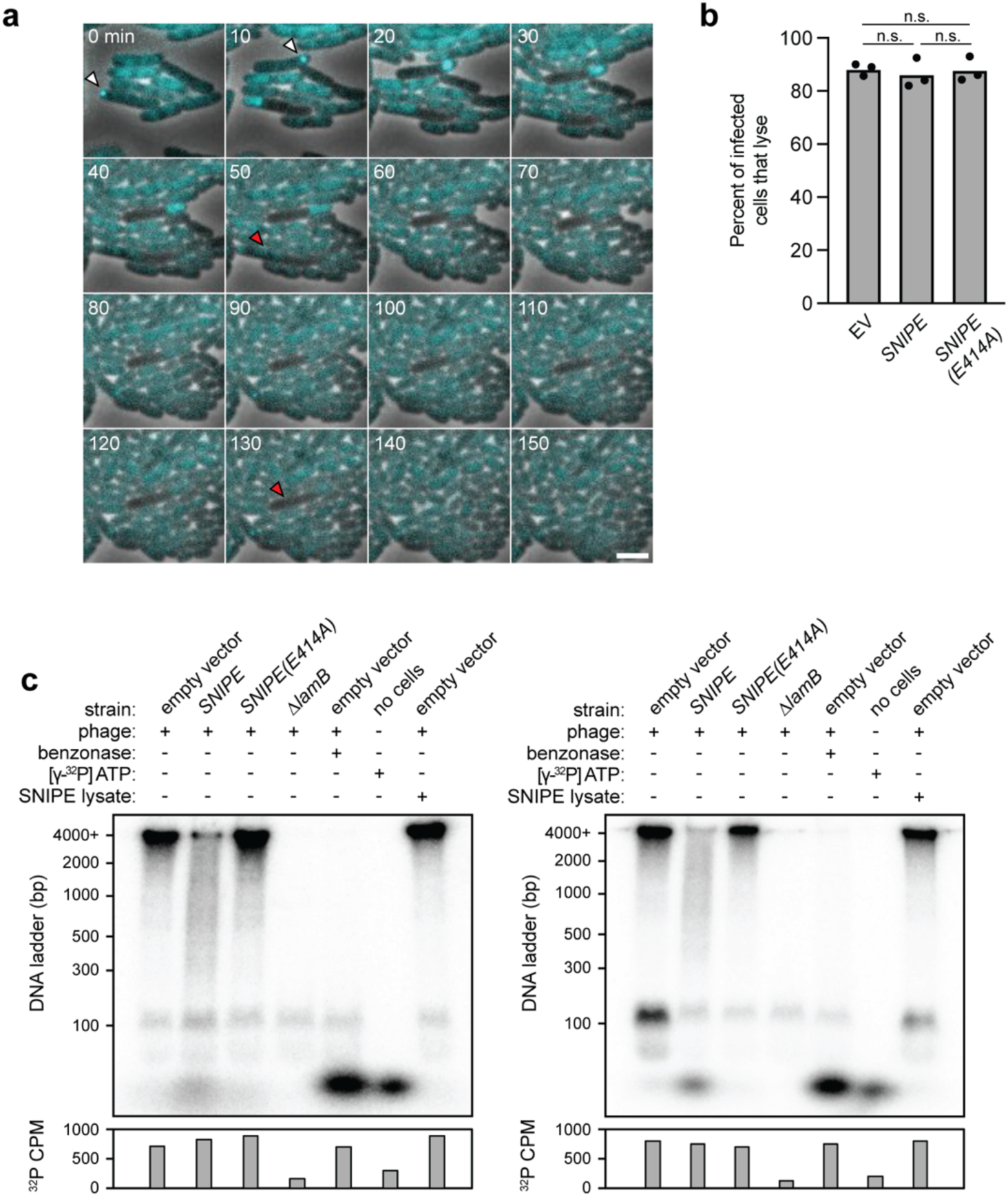
: Cell lysis of λ infection events that evade SNIPE and SNIPE-mediated cleavage of phage DNA. **(a)** Cells expressing SNIPE and CFP-ParB were infected with λ*^parS^*and imaged with time-lapse microscopy. The experiment is identical to Fig. 2b but a different field of view is shown for clarity in montage format. Phase and CFP channels are merged. White arrows indicate phage infection events. Red arrows indicate infected cells that lyse in the subsequent frame. Scale bar, 2 µm. **(b)** Cells which showed visible CFP-ParB foci during λ*^parS^* infection were tracked with time-lapse microscopy and the percent of these cells that lysed were quantified. Analysis was performed on samples shown in Fig. 2b-c and two additional independent replicates. n.s. indicates p > 0.5 (unpaired two-tailed t-tests). **(c)** Different bacterial strains were infected ^32^P-labeled λ and lysed 15 minutes after genome injection. Benzonase was added to one lysate sample. Lysates were subjected to electrophoresis through a polyacrylamide gel and imaged with a phosphorscreen. Total ^32^P per sample was measured with a scintillation counter. The phosphorscreen image on the left is identical to Fig. 2d but shows two additional controls. One control was [γ-^32^P] ATP in the absence of cell lysate and phage. In the other control, lysate from empty vector cells infected with ^32^P-labeled λ was mixed with uninfected SNIPE-expressing cell lysate to test if SNIPE can cleave phage DNA after lysis. An independent biological replicate is shown on the right. The bands at ∼50 and ∼100 bp were considered as nonspecific background due to their presence in the *ΔlamB* strain. For gel source data, see Supplementary Figure 1c.

**Extended Data Figure 4.**
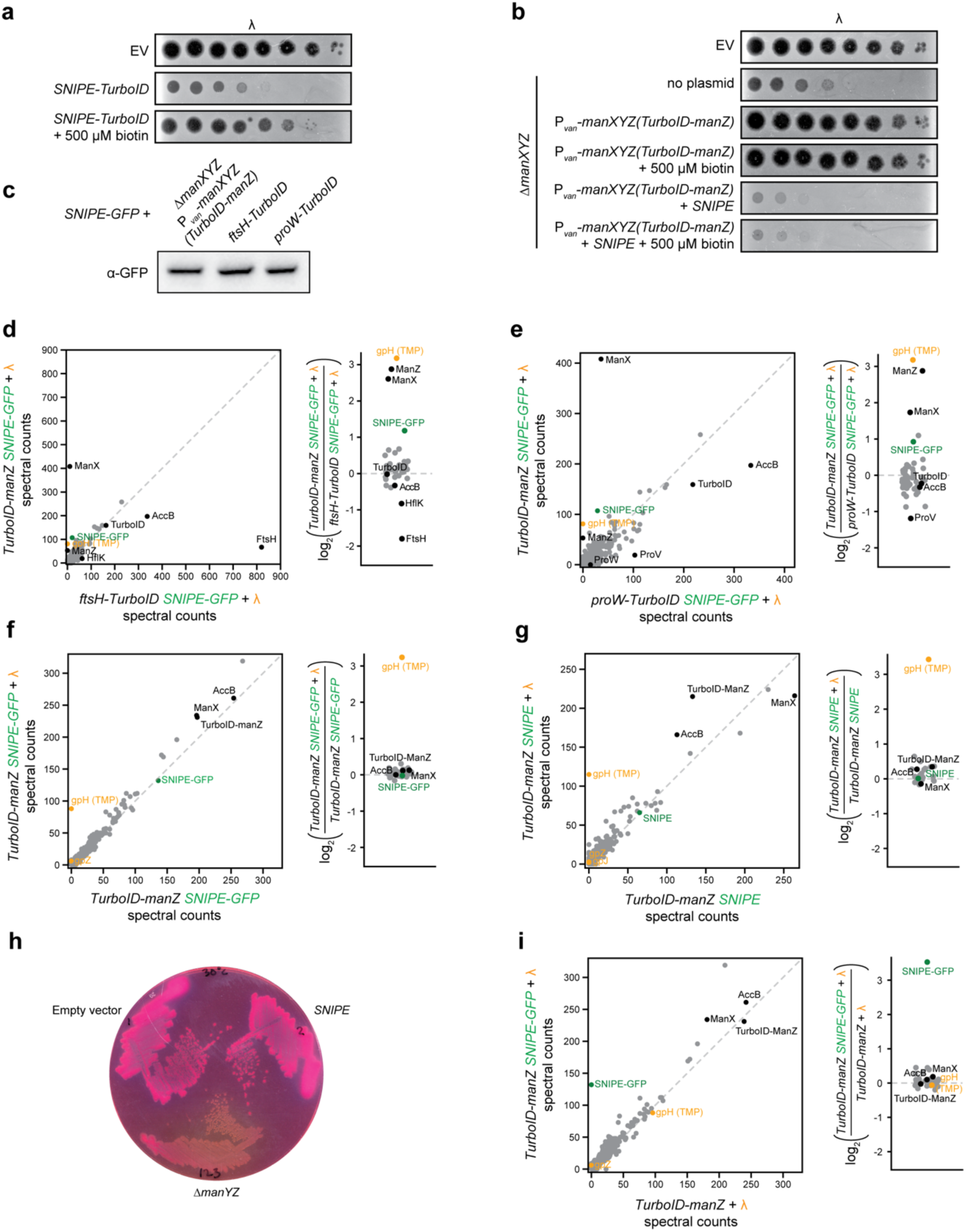
: Proximity labeling with TurboID. **(a)** Serial dilutions of λ were spotted onto lawns of cells expressing an empty vector or SNIPE-TurboID, in the presence or absence of 500 µM biotin. **(b)** Serial dilutions of λ were spotted onto lawns of cells expressing an empty vector or *ΔmanYZ* cells expressing no plasmid, TurboID-ManZ, and/or SNIPE, in the presence or absence of 500 µM biotin. **(c)** Expression levels of SNIPE-GFP in different TurboID fusion protein backgrounds, as assessed with immunoblots. The same membrane was stained with Coomassie as a loading control. For gel source data, see Supplementary Figure 1a. **(d)** Proximity labeling was performed with infected cells expressing SNIPE-GFP and TurboID-ManZ or FtsH-TurboID. Biotinylated proteins were enriched with streptavidin beads and quantified by mass spectrometry. Log_2_ ratios of spectral counts for proteins with ≥ 50 spectral counts in at least one sample are shown on the right. Phage proteins are labeled in orange, SNIPE is labeled in green, and other proteins of interest are labeled in black. Data is identical to Fig. 3c but zoomed out to permit visualization of all data points. **(e)** Same as **(d)**, but comparing infected cells expressing SNIPE-GFP and TurboID-ManZ or ProW-TurboID. **(f)** Same as **(d)**, but comparing infected versus uninfected cells expressing SNIPE-GFP and TurboID-ManZ. **(g)** Same as **(d)**, but comparing infected versus uninfected cells expressing SNIPE and TurboID-ManZ. **(h)** Empty vector, *SNIPE*, and *ΔmanYZ* colonies grown on MacConkey agar + 1% w/v mannose. Pink colonies indicate successful metabolism of mannose. **(i)** Same as **(d)**, but comparing infected cells expressing TurboID-ManZ versus cells expressing TurboID-ManZ and SNIPE-GFP.

**Extended Data Figure 5.**
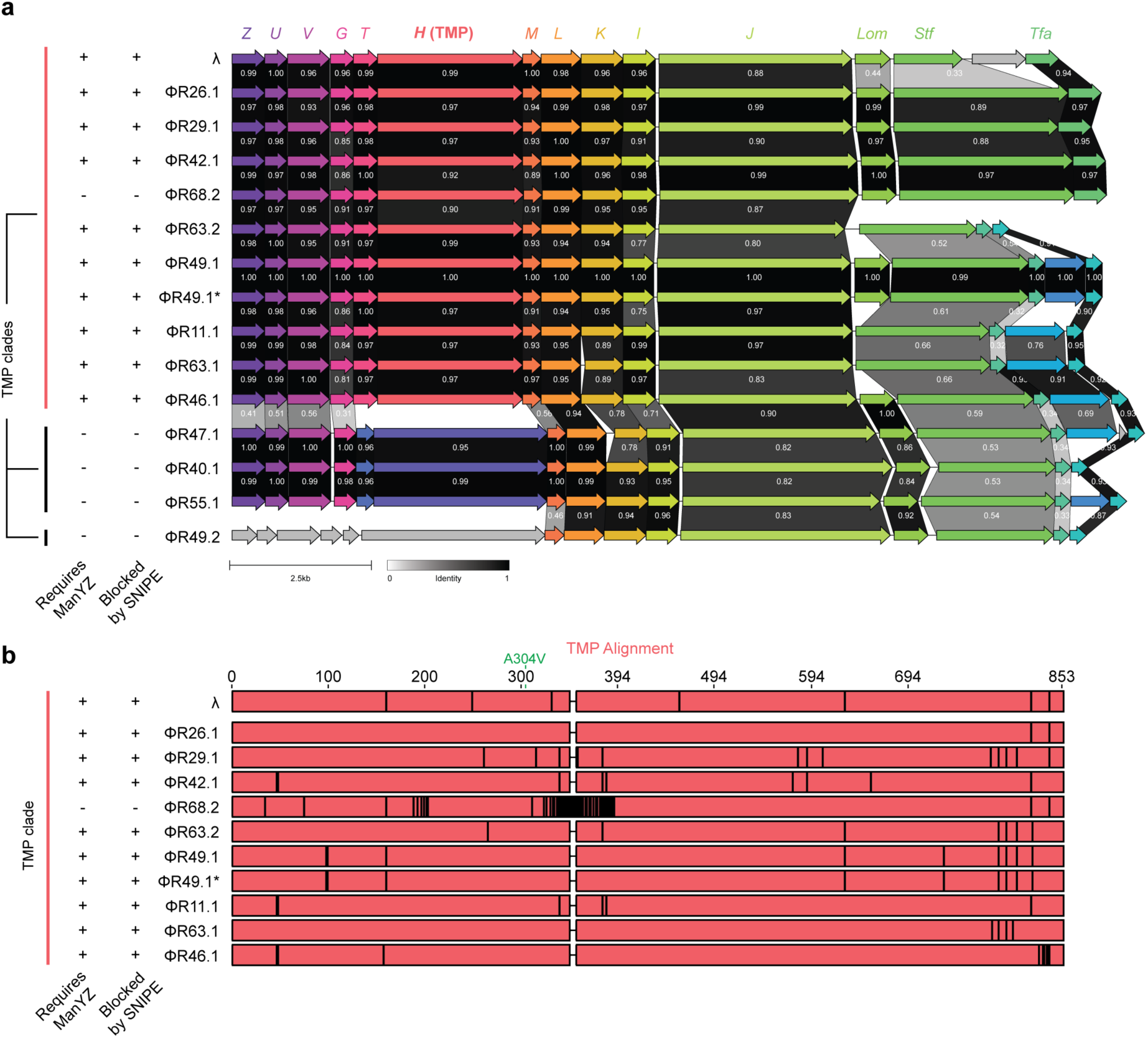
: **Alignments of tail genes from λ and a temperate phage panel. (a)** Tail gene clusters visualized with the CAGECAT clinker web server^67^. Sequence identity between genes (on a scale of 0 to 1) is shown. λ genes are labeled for reference. The propensity of each phage to be targeted by SNIPE and use ManYZ for genome entry is derived from data in Fig. 3f. Phages in a given TMP clade share > 90% amino acid identity between their TMPs, and < 20% identity with TMPs outside of the clade. **(b)** Alignment between TMPs of λ and other phages with the same TMP clade. Black lines indicate disagreements with the consensus sequence of the alignment. A304V indicates the mutation that is present in the generalist λ mutant, which allows it to circumvent a requirement for ManYZ. Amino acid numbers in the λ TMP are shown for reference. The propensity of each phage to be targeted by SNIPE and use ManYZ for genome entry is derived from data in Fig. 3f.

**Extended Data Figure 6.**
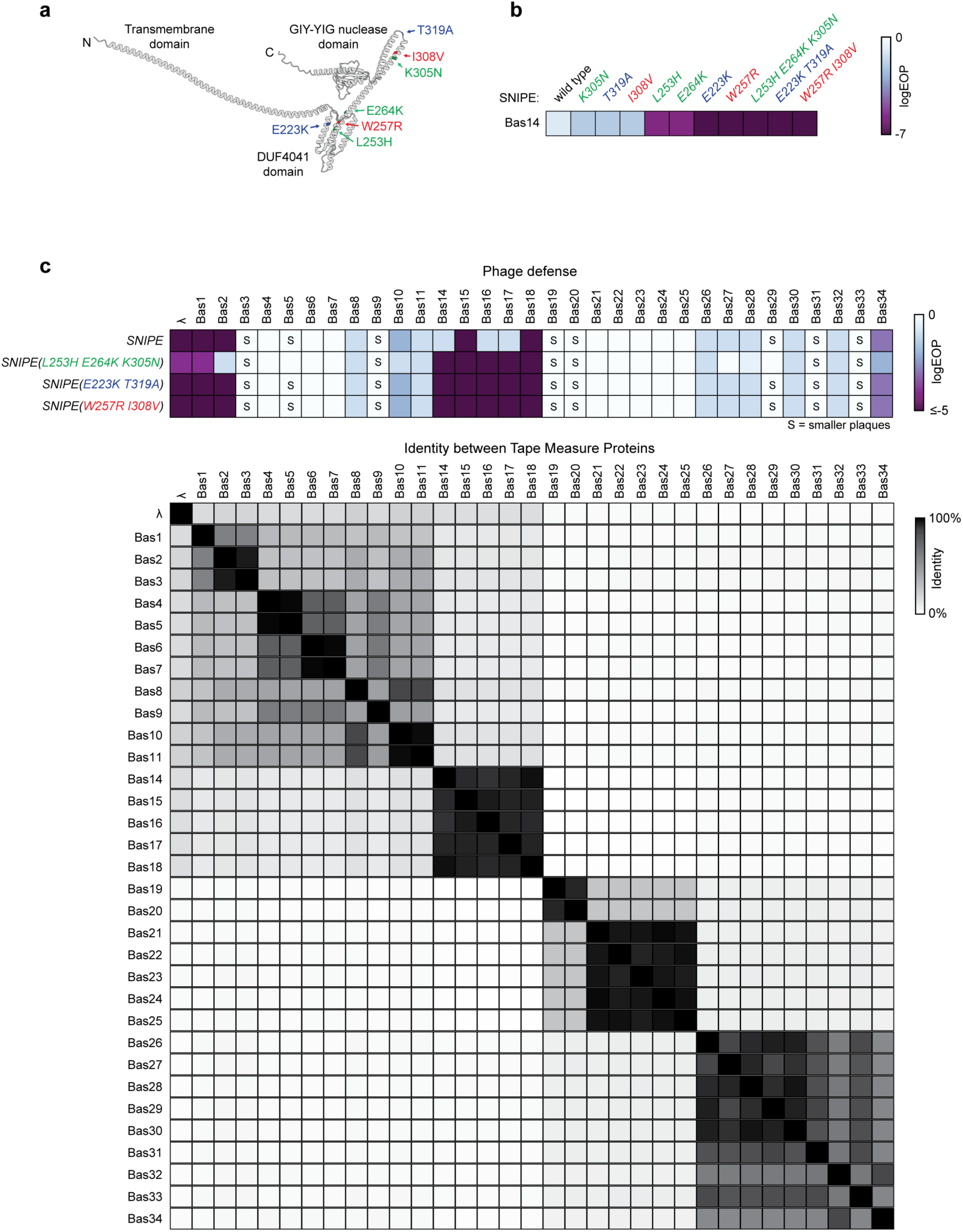
: SNIPE mutations enhance defense against the Bas14-18 clade. **(a)** Mutations that enhance SNIPE-mediated defense against Bas14 are highlighted on the structure of SNIPE predicted by AlphaFold3. The same schematic is shown in Fig. 4c and shown here for reference. **(b)** EOP data for Bas14 on SNIPE and different SNIPE mutants. **(c)** EOP data for λ and BASEL siphoviruses on SNIPE and different SNIPE mutants. Smaller plaque sizes are indicated with an ’S’. Amino acid sequence identity between TMPs of these phages is shown in matrix format.

**Extended Data Figure 7.**
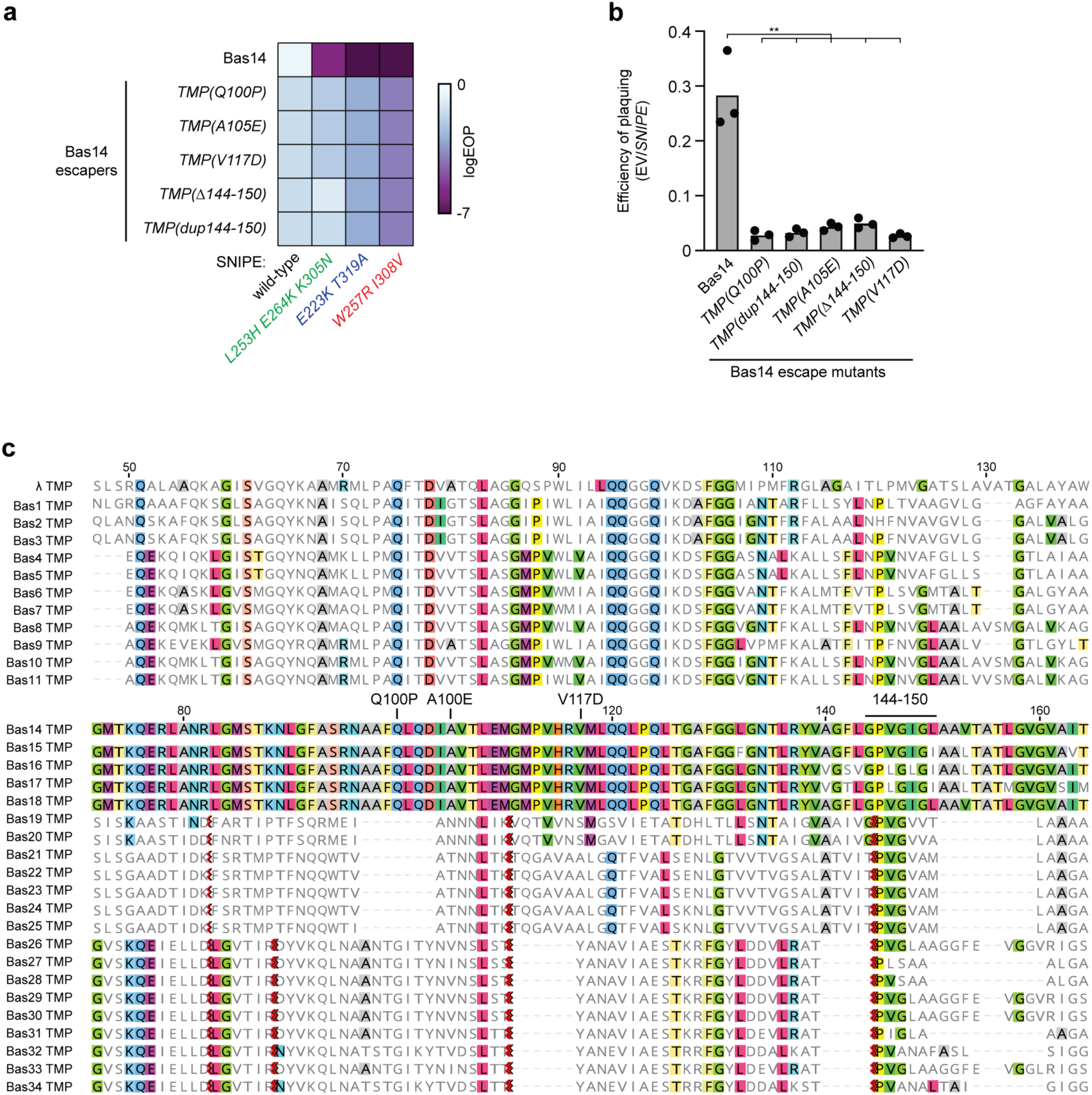
: Bas14 escape phenotypes and TMP alignment. **(a)** EOP data for Bas14 and Bas14 escape mutants on SNIPE and the indicated SNIPE mutants. **(b)** EOP data for Bas14 and Bas14 escape mutants on SNIPE. EOP was calculated by mixing phage and cells in top agar and using the top agar overlay method. Analysis was performed on three independent replicates. ** indicates p < 0.01 (unpaired two-tailed t-tests). **(c)** Alignment of the region of interest from λ and BASEL siphovirus TMPs. Locations of Bas14 escape mutations are shown above the Bas14 TMP sequence. Amino acid numbers in the λ TMP and Bas14 TMP are shown for reference. Amino acids highlighted in color indicate positions identical to the Bas14 TMP reference sequence, with specific colors corresponding to different residues. Red chevrons indicate gaps that were hidden.

**Extended Data Figure 8.**
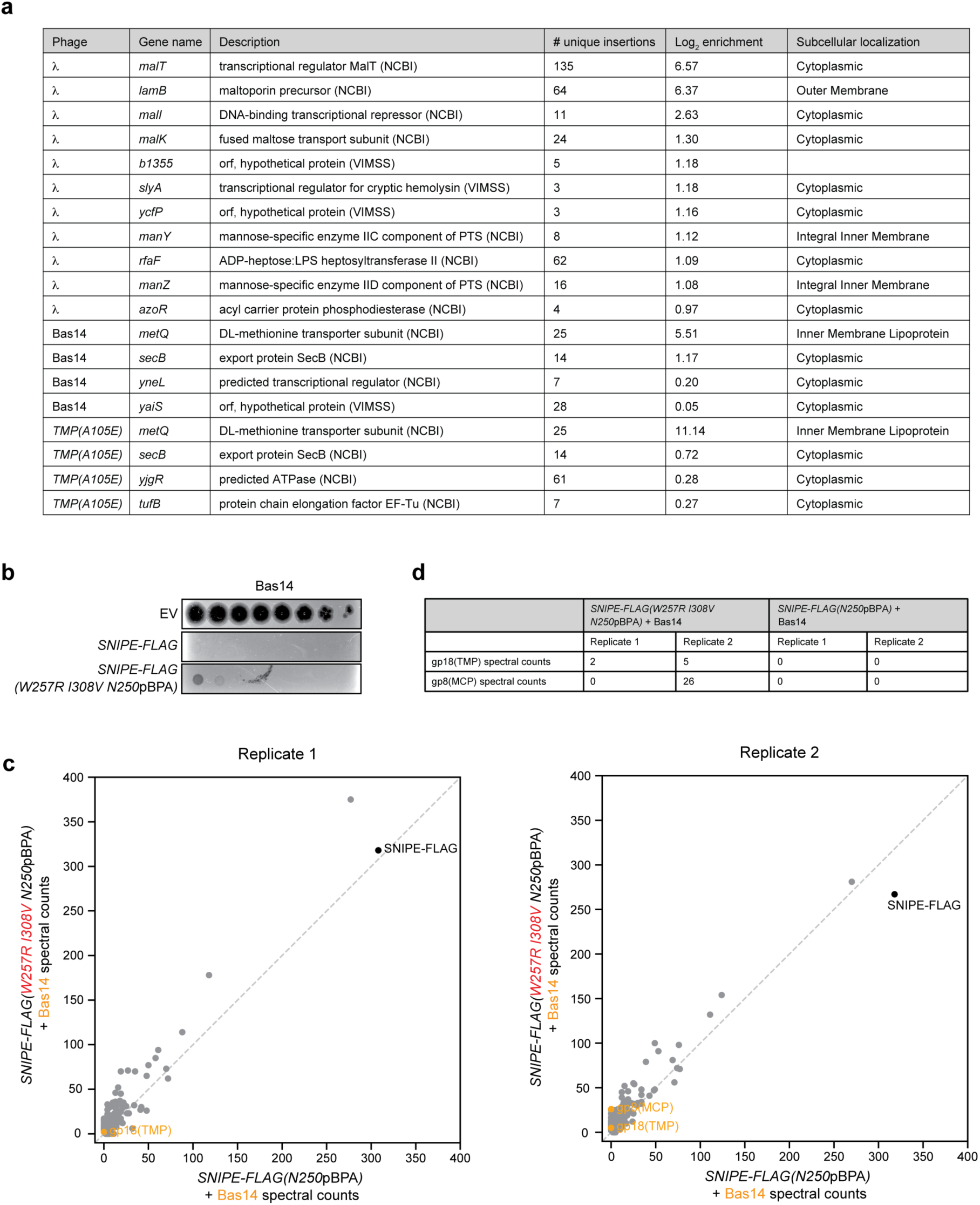
: Tn-Seq and pBPA crosslinking during phage infection. **(a)** List of host genes required for infection of λ, Bas14, and Bas14 *TMP(A105E)*, as identified via Tn-Seq. The number of unique transposon insertions per gene in the original pool, prior to selection, is indicated. Log_2_ enrichment scores refer to the averaged enrichment scores of transposons within a gene of interest after selection with a specific phage. Only genes with enrichment scores >0.95 are shown for λ, and ≥0.2 for Bas14 and Bas14 *TMP(A105E)*. The predicted subcellular localizations of gene products, as determined by UniProt^68^, are shown. **(b)** Bas14 phage spotting on strains expressing a specialized tRNA sythetase (pBpaRS) and an empty vector or different SNIPE-FLAG constructs with 1.2 mM pBPA. **(c)** UV-induced crosslinking of Bas14-infected cells expressing pBpaRS and SNIPE-FLAG(W257R I308V N250pBPA) or SNIPE-FLAG(N250pBPA) in the presence of 1.2 mM pBPA. These proteins and crosslinked products were pulled down with anti-FLAG beads and quantified by mass spectrometry. Phage proteins are labeled in orange, and SNIPE-FLAG is labeled in black. Two independent biological replicates are shown. Both replicates are identical to Fig. 4e but zoomed out to permit visualization of all data points. **(d)** Spectral counts of all phage proteins observed in **(c)**, shown in table format.

**Extended Data Figure 9.**
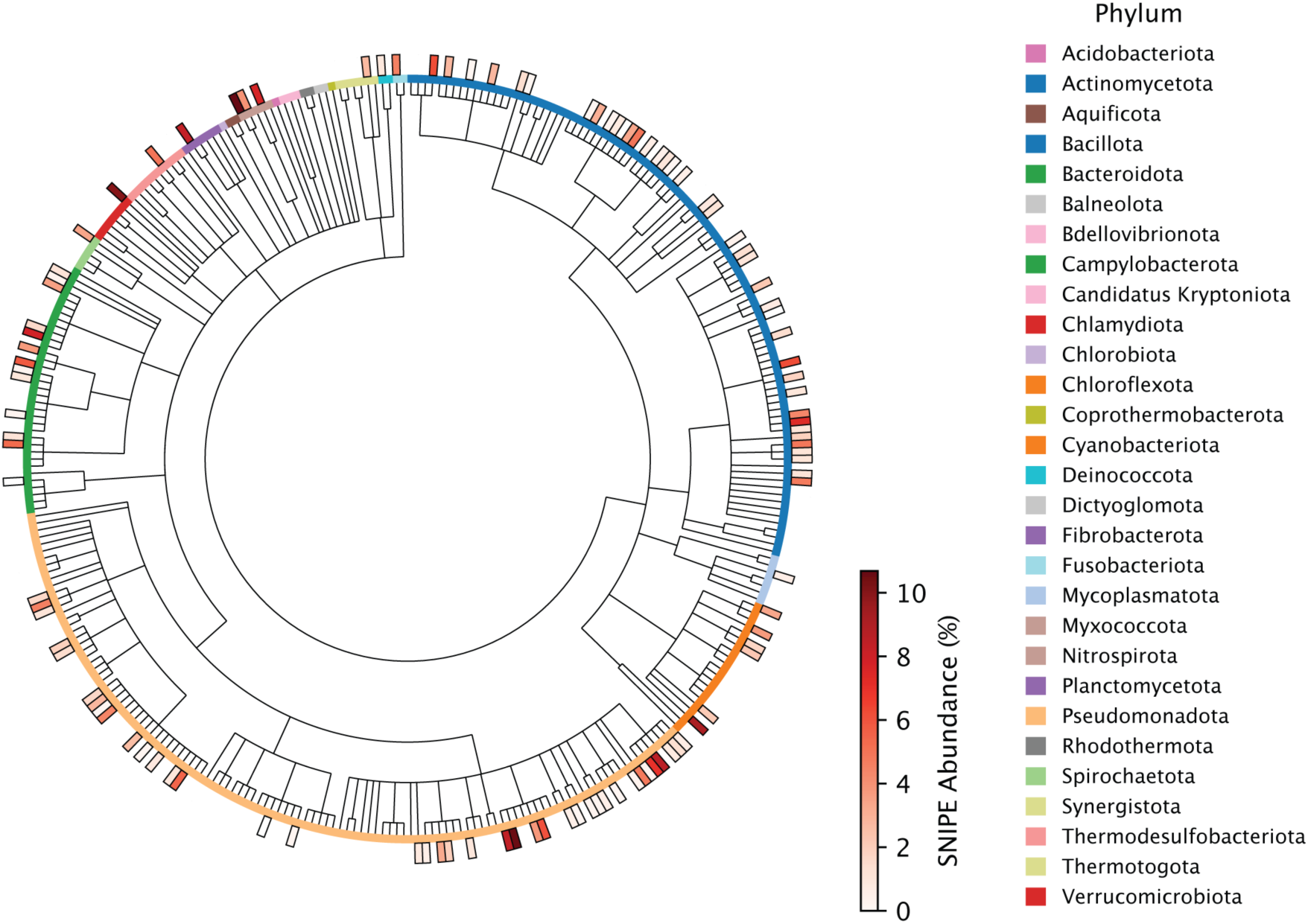
: Abundance of SNIPE across bacterial classes and phyla. Rooted phylogenetic tree of bacterial families, with phyla indicated by color in the inner ring. The outer ring indicates the percentage of genomes within each family that contain at least one SNIPE homolog. Black outlines indicate families that harbor at least one genome containing a SNIPE homolog. SNIPE homologs are as identified from Genbank and RefSeq genomes in a previous study^17^. Families are only shown if they contain ten or more genomes in the queried dataset.

**Extended Data Figure 10.**
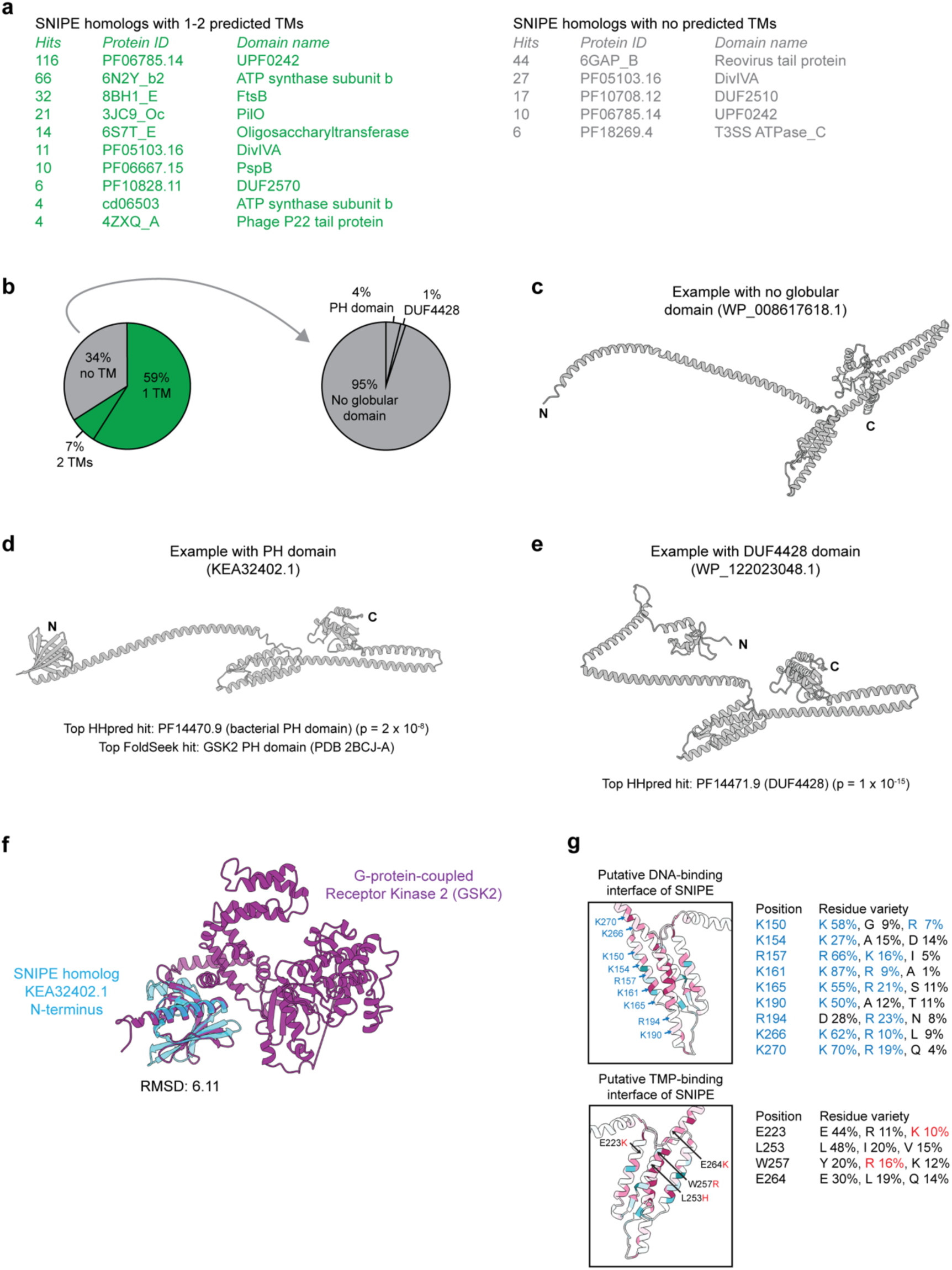
: Domains in N-terminal regions of SNIPE homologs. **(a)** Most frequent HMMER hits for the N-terminal regions of SNIPE homologs with 1-2 predicted TMs (left) or with no predicted TMs (right). Details of the hit count calculation are provided in the Methods section. **(b)** Structures of the 34% of SNIPE homologs that lack a predicted TM were predicted by AlphaFold2 and the percentages of homologs with a globular domain at their N-terminus (PH or DUF4428) are shown. Homology was detected by FoldSeek and/or HHpred. **(c)** Example of a predicted SNIPE homolog structure generated by AlphaFold2. The structure lacks both a predicted transmembrane (TM) domain and a globular domain in the N-terminal region. **(d)** Example of a predicted SNIPE homolog structure generated by AlphaFold2. The structure lacks a predicted transmembrane (TM) domain and features a predicted pleckstrin homology (PH) domain at the N-terminus. Top matches from FoldSeek (using the PDB100 database of experimentally determined structures) and an HHpred search with the N-terminal globular domain are presented. **(e)** Example of a predicted SNIPE homolog structure generated by AlphaFold2. The structure lacks a predicted transmembrane (TM) domain and features a predicted DUF4428 domain at the N-terminus. The top match from an HHpred search with the N-terminal globular domain is presented. **(f)** Structural alignment between the predicted PH domain of SNIPE homolog KEA32402.1 and the PH domain of GSK2 (PDB 2BCJ-A). **(g)** Same insets as shown in Fig. 5g, but with the percentage of different residues found at a given location across SNIPE homologs, as quantified by ConSurf. Only the top three most abundant residues are shown for each location. Positively charged amino acids are labeled in blue for the top inset. Mutations that enhanced defense against Bas14 are marked in red for the bottom inset.

## Supplemental Movie Legends

**Supplemental Movie 1:** Cells expressing SNIPE, PD-λ-3 (Abi system), or an empty vector (no defense) were infected with λ at a concentration such that half of the cells were infected at 0 minutes. Time-lapse imaging was performed with phase contrast microscopy for 180 minutes. Samples are identical to those shown in Fig. 1A. Scale bar, 3 µm.

**Supplemental Movie 2:** Microscopy of cells expressing CFP-ParB and an empty vector or different SNIPE constructs. Cells were infected with λ^parS^ and imaged with time-lapse microscopy for 150 minutes. The first column is phase contrast, the second column is CFP, and the third column is merged. Scale bar, 3 µm.

**Supplemental Movie 3:** λ*^parS^* lysogens expressing the heat-labile cI857 repressor, CFP-ParB, and SNIPE or an empty vector were induced via heat shock at 42 °C. Time-lapse microscopy was performed over 150 minutes to monitor CFP-ParB foci dynamics and cell lysis. The first column is phase contrast, the second column is CFP, and the third column is merged. Scale bar, 3 µm.

